# Modular engineered colorectal tumor microenvironments for immuno-oncology research and drug development

**DOI:** 10.64898/2026.06.02.729459

**Authors:** Adrian M. Filip, Barbora Lavickova, Aurelie Herault, Irineja Cubela, Benjamin Ehret, Esther Danenberg, Michelle Daniel, Stéphane Chevrier, Mairene Coto-Llerena, Giulia Protti, Soren Muller, Sophie Jansen, Kristina Kromer, Marius F. Harter, Sofia Tsiepa, Ilya Lukonin, Wenjie Jin, Pol Jean-Mairet, Floriana Cremasco, Sara Colombetti, Lauriane Cabon, Nikolche Gjorevski

**Author notes:** these authors contributed equally to this work (Adrian M. Filip, Barbora Lavickova).

## Abstract

The development of cancer immunotherapies is hindered by the lack of human-relevant models that accurately translate to patient outcomes. We combine patient-derived colorectal tumor organoids (PDOs) and cancer-associated fibroblasts (CAFs) into floating extracellular matrix drops to form miniature colorectal tumors. These SHaking Organoid CO-cultures (SHOCOs) reproducibly capture key features of the tumor microenvironment (TME), including physiologically relevant ECM, hypoxic regions, and a diversified stromal compartment that captures primary CAF states. Compared with traditional PDO-based co-culture models, SHOCOs maintain immune cells in numbers, states and functional interactions that more closely reflect the physiological tumor microenvironment. The modularity of the system allowed controlled simulation of diverse patient tumor archetypes, and defining how TME components shape therapy response. Immune-rich SHOCOs treated with T-cell bispecific antibodies exhibited robust anti-tumor responses that were stronger and more rapid than their PDO-based counterparts. Stroma-rich tumors, on the other hand, diminished therapy responses by physically hindering both immune cell recruitment and limiting antibody penetration. Thus, SHOCOs enable the construction of modular tumor microenvironments, providing a versatile platform to dissect immune and stromal tumor biology and to catalyze the discovery of novel therapeutic approaches.

## Introduction

The immense success of immunotherapies like immune-checkpoint inhibitors (ICIs) and chimeric antigen receptor T cells (CAR-T) has spurred widespread efforts to leverage immune cells in battling cancer, thus igniting hope for a broad and durable cure^1–4^. However, after the initial leaps, progress has been modest, with novel drugs experiencing an attrition rate of 95% in Phase I clinical trials currently^5,6^. The high failure rate unambiguously signals the poor predictive value of immuno-oncology models, suggesting that compelling results from basic research and preclinical drug development do not translate to patients^7^.

Widely used mouse models and human cell-based culture systems must be credited for their role in the success of ICIs and CAR-T therapies. However, slow progress in targeting solid, epithelial tumors has highlighted their limitations. In particular, human culture systems lack the multifactorial, spatial and temporal complexity of the tumor microenvironment (TME)^8,9^, whereas mouse models feature a TME with sufficient complexity but questionable fidelity to actual human tumors^10–12^. To address the latter, efforts are underway to create sophisticated, fully humanized mouse models which will inevitably catalyze the discovery of novel and effective immunotherapies^13^.

In parallel, researchers have begun creating complex and human-relevant TME models *in vitro*. Patient-derived organoids (PDOs) are faithful representations of the tumor epithelium, capturing its mutational landscape, morphological features and response to epithelium-targeted drugs^14–18^. To introduce other TME components, PDOs have been co-cultured with diverse immune and stromal cells^19–27^, and even whole tumor digests^28^. Collectively, these models capture increasingly sophisticated aspects of the TME, yet often excel along different axes of complexity, scalability and experimental control. An outstanding challenge is therefore to unite these strengths in a single system that combines reproducibility, modularity and scalability with the structural and biological complexity of native tumors^29–32^.

We combined colorectal cancer (CRC) PDOs, tumor-relevant extracellular matrix (ECM), cancer-associated fibroblasts (CAFs) and immune cells in shaking co-culture to create human-relevant models of CRC, which we term SHOCOs (SHaking Organoid CO-cultures) (**Figure 0**). Owing to the contractile activity of CAFs and the mechanical properties of the ECM, SHOCOs feature a dense, tissue-like internal structure, and exhibit regionalization akin to that of native tumors. Exploiting the control afforded by the model, we generated immunologically “hot”, “cold” and stroma-dense TMEs, and examined how they influence immunotherapy responses. Immune-rich SHOCOs displayed robust concentration- and time-dependent responses to T-cell bispecific antibodies (TCBs) that were faster and stronger than those observed in conventional PDO-based co-cultures. Stroma-rich SHOCOs undercut anti-tumor activity by halting T-cell infiltration, as well as TCB penetration and target engagement. Together, these findings establish SHOCOs as a scalable and versatile platform for constructing defined tumor microenvironments and systematically interrogating how their composition shapes tumor biology and therapeutic responses.

**Figure 0.**
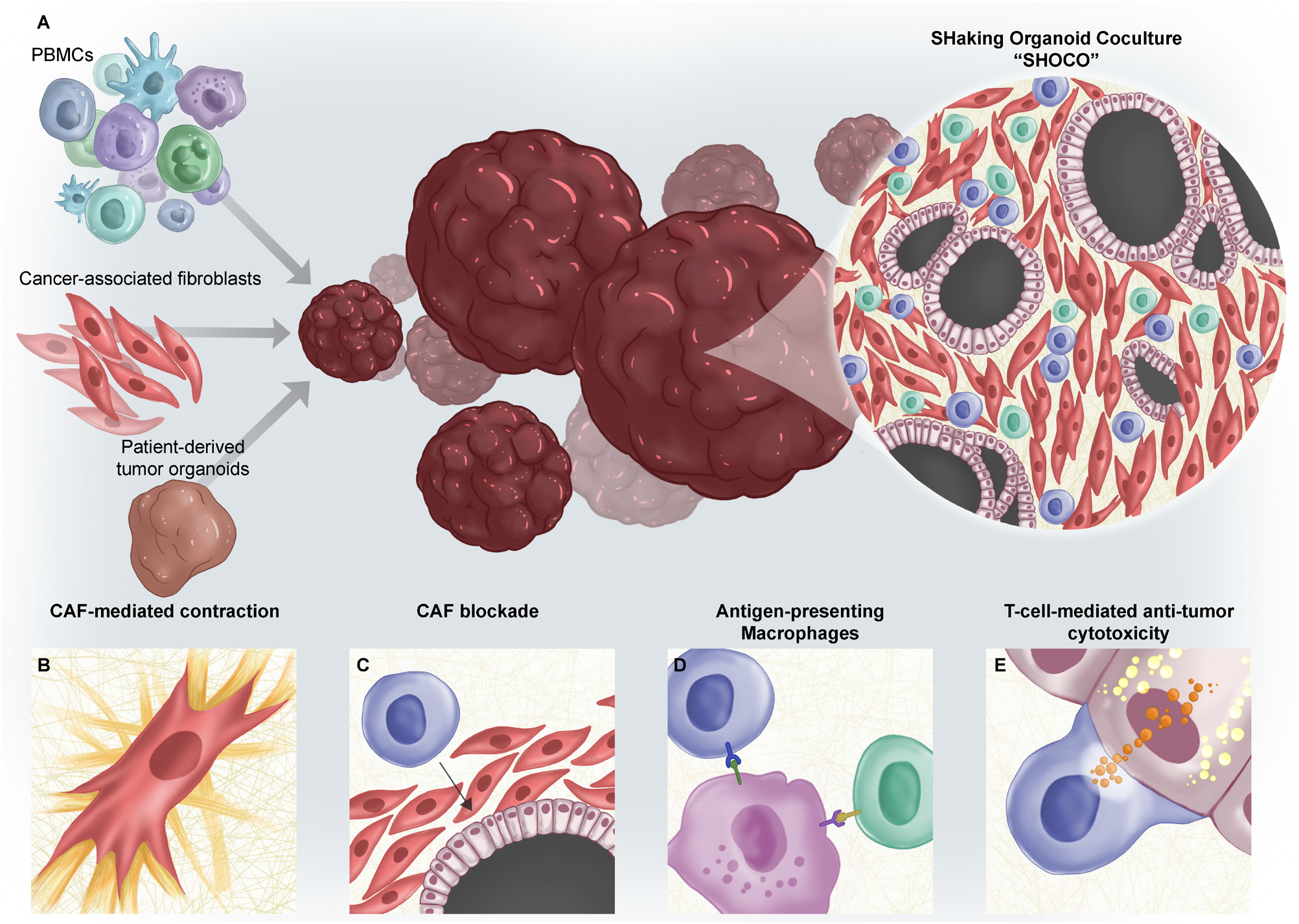
Graphical Abstract: Illustration of the SHOCO Model. **(A)** Overview of the components used in creating SHOCOs and the complex tumor microenvironment they replicate. Epithelial cells (pink), CAFs (red), CD8 T cell (blue), CD4 T cell (green). Key SHOCO characteristics highlighted by: **(B)** Emphasizing the role of CAFs in contracting and creating a denser microenvironment. **(C)** CAFs acting as a physical barrier, preventing CD8 T cells from reaching tumor niches. **(D)** SHOCOs maintaining a substantial myeloid population (violet) capable of performing antigen presentation. **(E)** T cells within SHOCOs mounting an anti-tumor cytotoxic immune response as a result of immunotherapy. Granzymes (orange), cleaved Caspase-3/7 (yellow).

## Results

### PDOs and autologous CAFs in shaking co-culture recreate the heterogeneous milieu of primary colorectal tumors

To establish compact 3D structures that feature a densely packed stroma reminiscent of that occurring within native colorectal tumors (**Figure S1A**), we provided a mechanically unconstrained environment that is amenable to CAF-mediated tissue remodeling. More specifically, we generated donor-matched PDOs and CAFs from CRC resections and encapsulated them in a floating droplet, formed via surfactant-stabilized water-in-oil emulsion (**Figure 1A**). The aqueous phase housing the TME components comprised type I collagen or reconstituted basement membrane hydrogel precursors, which polymerized into a solid gel. After gelation, the structures were transferred to a shaking culture. Owing to CAF-mediated contraction of the extracellular matrix (ECM) mesh, the floating droplets were compacted into dense mini-tumors, which we term SHOCOs (SHaking Organoid CO-culture). Importantly, the simplicity of this process allowed for automated and large-scale production of SHOCOs, simultaneously increasing their quality and reproducibility.

**Figure 1.**
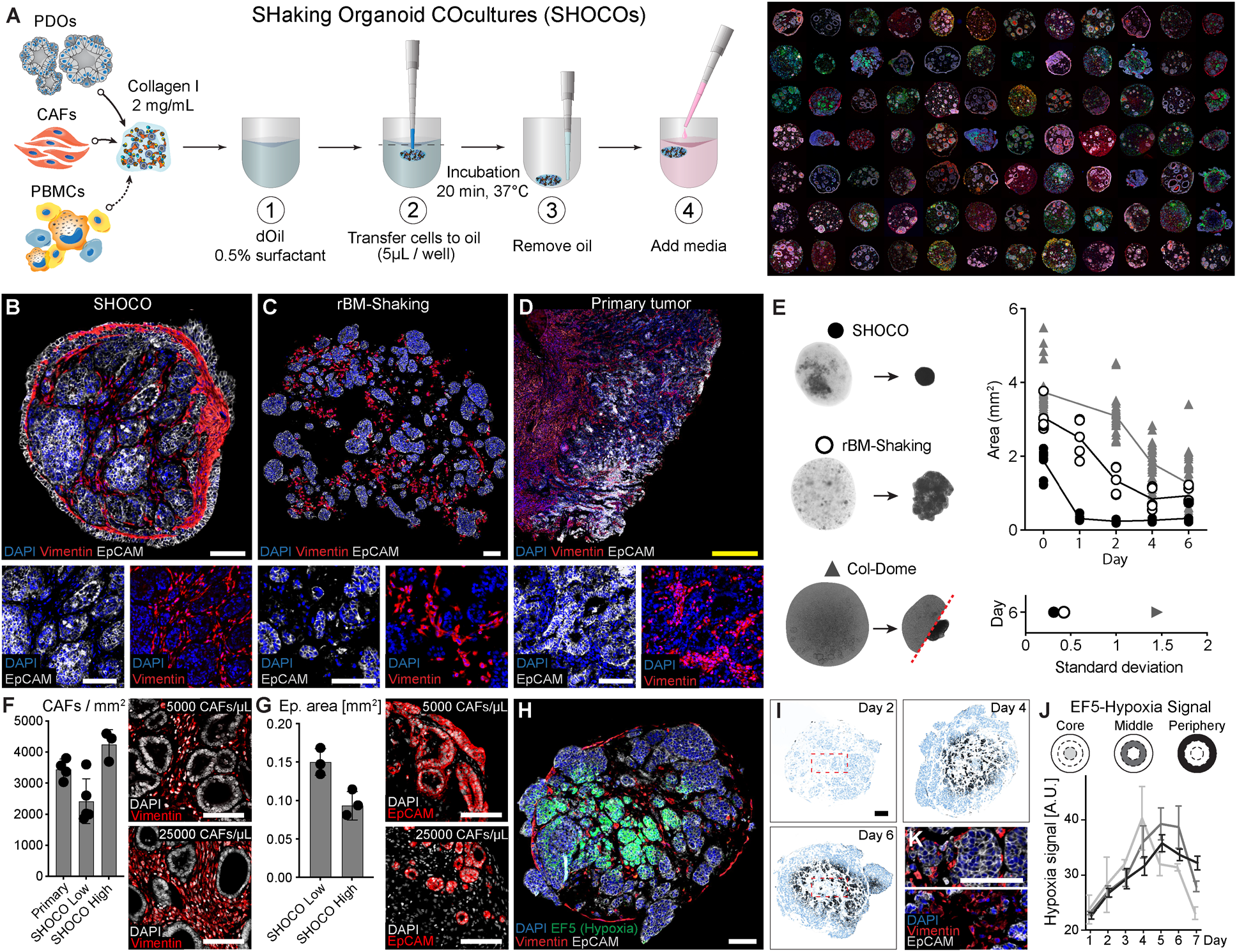
Patient-derived intestinal organoids and matched cancer-associated fibroblasts co-cultured in a shaking collagen-rich matrix consistently recreate key architectural and microenvironmental features of primary tumors. **(A)** Schematic of the high-throughput shaking organoid co-culture (SHOCO) workflow integrating patient-derived tumor organoids (PDOs), matched cancer-associated fibroblasts (CAFs), and peripheral blood mononuclear cells (PBMCs), with a montage of representative SHOCOs. **(B–D)** Representative multiplex immunofluorescence (mIF) images of a collagen-based SHOCO (B), rBM-based SHOCO (rBM-Shaking) (C), and corresponding primary tumor (D) from Donor 5. EpCAM and vimentin mark epithelial and stromal compartments, respectively. **(E)** Quantification of projected area of collagen-based SHOCOs, rBM-Shaking cultures, and Col-Dome cultures over 6 days. The lower panel shows the standard deviation of day-6 area for each culture condition. *N* = 2 donors (Donor 1/Donor 4); *n* = 5/4 SHOCOs, 2/3 rBM-Shaking cultures and 15/16 Col-Dome domes. **(F)** Quantification of CAF density in primary tumors and SHOCOs generated at low (5000 CAFs/µL) or high (25000 CAFs/µL) CAF density, with accompanying vimentin mIF images. Primary tumor: *N* = 3 (Donor 9 / Donor 10 / Donor 11); *n* = 2/1/1 stromal regions per donor. SHOCOs: *N* = 4 donors across two experiments (Donor 1, Donor 4, Donor 5, Donor 6); *n* = 5 donor-experiment means from two experiments. Donor 5 was included in both experiments. Means comprised approximately 10 SHOCOs (136 FFPE images total) at 5000 CAFs and 7 SHOCOs (84 FFPE images total) at 25000 CAFs. Individual replicates and ranges are shown. **(G)** Quantification of epithelial niche area in SHOCOs generated at low or high stromal density, with accompanying EpCAM mIF images. *N* = 3 donors (Donor 1 / Donor 4 / Donor 5); *n* = 3 donor means/condition. Each dot is a donor-specific mean comprising approximately 8 SHOCOs/donor (92 FFPE images total) at 5000 CAFs and 7 SHOCOs/donor (84 FFPE images total) at 25000 CAFs. Individual replicates and ranges are shown. **(H)** EF5 staining of a day-6 SHOCO (Donor 7) showing spatially localized hypoxic regions. **(I)** Spatial distribution of EF5-positive regions in SHOCOs at days 2, 4, and 6. **(J)** Quantification of EF5 signal across 100 μm concentric regions defining the core, middle, and periphery of SHOCOs over 7 days. *N* = 1 (Donor 7); *n* = 3 SHOCOs per timepoint; 3 FFPE images per SHOCO. Data are presented as mean ± SD. **(K)** High-magnification mIF images of SHOCO cores at days 2 and 6 showing EpCAM-positive epithelial and vimentin-positive stromal compartments. Scale bars, 100 μm (white and black) and 500 μm (yellow).

Next, we examined how the choice of ECM influences SHOCO structure and its concordance to primary colorectal tumors. Previous studies have used reconstituted basement membrane (rBM) hydrogels to form similar co-cultures that approximate the bladder, termed ‘bladder assembloids’^19^. Bearing in mind the collagen-rich interstitial matrix of most solid tumors^33,34^, we deemed type I collagen gels a more physiologically appropriate alternative. Comparing collagen- (**Figure 1B**) and rBM-based (**Figure 1C**) SHOCOs, we found that both systems featured a heterogeneous microenvironment, with tumor niches surrounded by a dense, stroma-like fibroblast mesh that mirrors their *in vivo* counterpart (**Figure 1D**). The tumor epithelium displayed the expected architecture and polarization^37,38^, including luminal enrichment of carcinoembryonic antigen-related cell adhesion molecule 5 (CEACAM5) (**Figure S2A**).

However, the two systems differed markedly in ECM organization and turnover (**Figures S1A** and **S1B**). Collagen-based SHOCOs developed structured ECM deposits, particularly along the tissue periphery, and actively reshaped their matrix over time, progressively accumulating collagens I, IV and V. In contrast, ECM proteins in rBM cultures remained dispersed around small CAF clusters and changed comparatively little over the same period. Fibronectin fibrils were virtually absent from rBM cultures but became widespread throughout collagen-based SHOCOs within 24 hours, consistent with the established role of fibronectin in fibroblast-mediated collagen contraction^35,36^. Collagen III was likewise detected only in collagen-based SHOCOs, although, given its strong day 0 signal, its expression cannot be attributed solely to *de novo* CAF secretion and reflects collagen III already present in the hydrogel at model formation (**Figures S1A** and **S1B**).

Differences in stromal organization within the two models were associated with distinct compaction dynamics. Collagen-based SHOCOs contracted almost immediately and reached their minimal size by day 2, whereas rBM cultures compacted gradually and remained consistently larger despite identical cellular composition (**Figures 1E** and **S2B**). The extent of CAF-mediated SHOCO contraction depended upon multiple parameters, including collagen abundance, donor differences and epithelial (PDO) influences (**Figure S2D**). In addition to comparing SHOCOs with collagen- and rBM-based ECM, we also compared collagen-based SHOCOs with compositionally matched PDO-CAF dome co-cultures (Col-Dome). We generally found working with Col-Domes to be challenging, because, owing to CAF mechanical activity, they frequently detached from the cell culture substratum and formed SHOCO-like structures. Unlike the floating cultures, however, this process was unpredictable and variable across donors and even among replicate domes from the same donor (**Figures 1E** and **S2C**). Thus, based on their technical robustness and higher structural fidelity to primary tumor resections, we deem collagen-based SHOCOs the most relevant CRC model among those tested. They form the main focus of the rest of the study, where we refer to them simply as ‘SHOCOs’.

Stromal abundance varies considerably across colorectal tumors and even between regions of the same tumor^39,40^. Given the presence of CAFs within our model, we next asked whether SHOCOs could be adapted to reproduce distinct stromal environments. By increasing the starting CAF concentration from 5000 cells/µL (CAF-low SHOCO) to 25000 cells/µL (CAF-high SHOCO), we generated SHOCOs with stromal densities falling below and above those measured in our primary colorectal tumor samples (**Figure 1F**). To quantify this difference, FFPE sections were segmented into epithelial and stromal regions and *a*-SMA^+^ cells were counted within the latter. Interestingly, increasing the stromal content came at a clear cost to the epithelial compartment: PDO niches were markedly smaller in CAF-high SHOCOs, indicating that epithelial expansion becomes increasingly constrained as the surrounding stroma grows denser (**Figure 1G**). Thus, simply varying the initial number of CAFs allowed us to generate stroma-poor and stroma-rich microenvironments, which can be used to investigate how stromal abundance influences immune behavior and response to therapy.

The presence of a hypoxic and, sometimes, necrotic core is a hallmark of the TME which governs a range of tumor behaviors and outcomes, including the anti-tumor immune response^41–46^. We hypothesized that the dense, fibrotic structure of the SHOCOs, along with their relatively large size (diameter of around 1 mm after contraction) would reproducibly create hypoxic zones, which are poorly captured in individual PDOs, owing to their small size, or conventionally used rBM-based co-cultures (rBM-Dome), owing to their low density and sparse cellular contacts. Indeed, EF5 staining revealed robust and well-defined hypoxic regions that emerged centrally and progressively extended toward the periphery (**Figures 1H-K**). The EF5 signal peaked in the center during the first 4 days and subsequently shifted toward the middle layer. Spatial mass cytometry supported this organization, with CAIX and GLUT1 closely following the hypoxic region (**Figure S2E**). As the hypoxic region progressed outward, epithelial and stromal structures were progressively lost from the center. TUNEL staining of serial sections from the same SHOCOs confirmed extensive cell death within this region (**Figure S2F**), further supporting the emergence of a necrotic-like core. Thus, beyond reproducing key cellular and stromal constituents, the size and density of SHOCOs give rise to spatial metabolic and cellular features characteristic of the native TME.

### SHOCOs promote diversification of *in vitro*-expanded CAFs into CRC-like stromal states

Next, we used single-cell RNA sequencing (sc-RNAseq) to characterize CAF states and diversity in SHOCOs, and benchmarked them against the starting 2D culture and conventional rBM-based PDO-CAF dome (rBM-Dome) co-culture (**Figure 2A**). 2D-cultured CAFs clustered separately from those incorporated into 3D systems, and chiefly inhabited myofibroblast-like (*ACTG2*, *TAGLN*, *MYL9*) and proliferative (*MKI67*, *TOP2A*) states, as reported extensively across the literature^47,48^ (**Figures 2B-D** and **S3A-C**). Incorporation into either 3D format led to a significant diversification of CAFs (**Figures 2B-D**), underpinned by the appearance of stromal subsets encoding for a wider range of ECM genes beyond *COL1A1* (*COL4A5*, *LAMA2, POSTN*), as well as ECM-binding integrins (*ITGA1*) (**Figures 2B-D** and **S3A-C**). This broader CAF repertoire is consistent with human CRC models showing that three-dimensional tumor co-culture can induce fibroblast states underrepresented in monoculture^49^. Comparing the two 3D models, SHOCOs supported the emergence of subsets with signatures related to matrix deposition and remodeling (*POSTN*, *MMP14*, *TIMP1*) and epithelial signaling (*NRG1*), which are essential for shaping various aspects of CRC progression^50–52^. Importantly, CAFs associated with ECM production and remodelling are known to promote stromal fibrosis and immune exclusion in solid tumors^53^. *POSTN*, which is uniquely upregulated in SHOCOs (**Figure S3D**) was specifically linked to stroma-mediated blockade of therapeutic antibody penetration in patient tumors^54^. Pathway analysis confirmed the proliferative state of 2D-cultured CAFs, and, within the 3D models, revealed an induction of the WNT and TGF- β pathways (**Figure 2E**) – master regulators of intestinal biology and CRC^54,55^. Compared with those in conventional co-culture, SHOCO CAFs were enriched in programs related to hypoxia and matrix remodelling, which is in line with our immunofluorescence data (**Figures 1H**, **S1** and **S2E-F**) and the SHOCO-specific induction of CAF subsets expressing genes that encode for ECM, ECM-binding integrins and MMPs (**Figures 2B-D**). Strikingly, SHOCOs appeared to uniquely harbor inflammatory signaling, exemplified by the TNFα–NF-κB pathway (**Figure 2E**).

**Figure 2.**
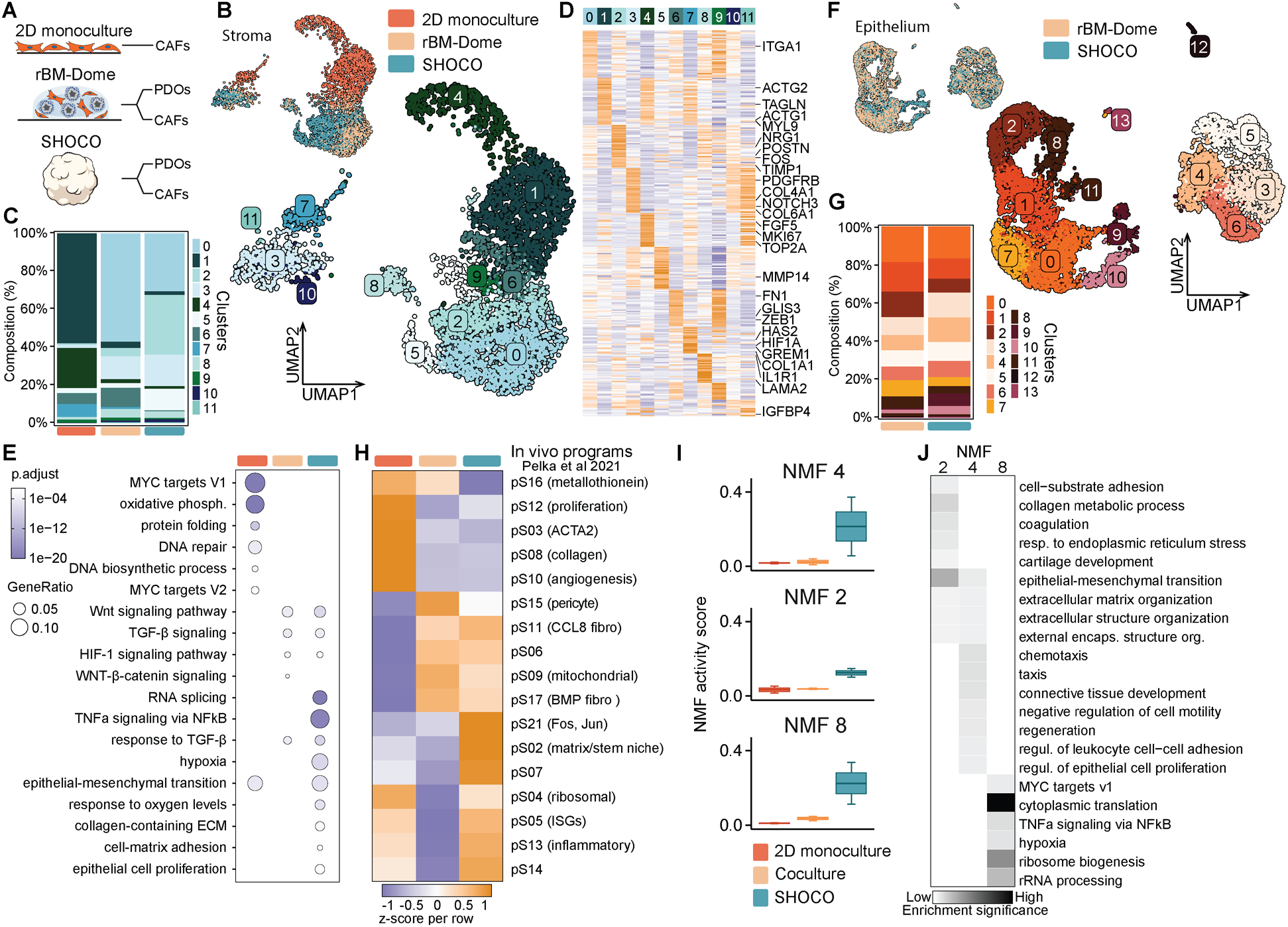
SHOCOs promote broad diversification of 2D-expanded CAFs into CRC-like stromal states while preferentially enriching matrix-remodeling, hypoxia-responsive and inflammatory programs. **(A)** Schematic of the culture systems used for scRNA-seq analysis: CAF 2D monoculture, rBM-Dome culture (PDO-CAF), and SHOCOs (PDO-CAF). **(B)** Integrated UMAP embedding of scRNA-seq transcriptome data from stromal cells isolated from CAF monoculture, from rBM-Dome cultures, or from SHOCOs. *N* = 2 (Donor 1, Donor 8). **(C)** Stacked bar plot showing the relative proportions of stromal cell subclusters within each culture condition. **(D)** Heatmap showing the expression patterns of marker genes across RNA clusters (Table S1). **(E)** Enrichment analysis (GO, KEGG, and MSigDB Hallmarks) for genes upregulated in stromal cells under different culture conditions (Table S1). Displayed pathways were manually selected for relevance. **(F)** Integrated UMAP embedding of scRNA-seq transcriptome data from epithelial cells isolated from rBM-Dome cultures or SHOCOs. *N* = 2 (Donor 1, Donor 8). **(G)** Stacked bar plot showing the relative proportions of epithelial cell subclusters within each culture condition. **(H)** Mapping of *in vivo* CRC fibroblast gene programs (Pelka et al., Table S1) across culture models. **(I)** Comparison of activity scores for cNMF-derived stromal gene-expression programs across culture systems enriched in SHOCO co-culture (additional programs are shown in **Figure S3H**). Data are shown as box-and-whisker plots with the median as the center line. **(J)** Pathway enrichment (GO Biological Process and MSigDB Hallmark) for the SHOCO-enriched cNMF program analyzed using clusterProfiler (Table S1).

By comparison, culture format had only a modest effect on the overall transcriptional landscape of the tumor epithelium, with SHOCO- and rBM-dome-derived cells showing similar cluster distributions (**Figures 2F** and **2G**). The notable exceptions were clusters 9 and 10, which were expanded in SHOCOs (**Figures 2F, 2G** and **S3E**). Cluster 9 was enriched for Integrated Stress Response and epithelial plasticity genes (*ANXA1*, *ANXA5*, *DDIT3*, *HSPA1A*), consistent with pathway enrichment in SHOCOs highlighting TNFα signaling via NF-κB, hypoxia, and inflammatory responses. Meanwhile, cluster 10 was enriched for a structural plasticity program (*MACF1*, *TRIO*, *PARD3*, *EXT1*) driving cytoskeletal remodeling and cell motility, corresponding to structural and migratory pathways including ECM organization, intermediate filament organization, and positive regulation of cell migration (**Figures S3F** and **S3G**).

To assess the correspondence of CAFs across the different *in vitro* models to those within primary CRC, we mapped stromal programs identified by Pelka et al.^31^ in patient samples to our scRNAseq dataset (**Figure 2H**). Overall, both 3D cultures captured patient stromal signatures, with SHOCOs spanning a broader range of these states. As expected, 2D CAFs displayed predominantly myofibroblast, proliferative and collagen-associated programs, reinforcing the contractile and biosynthetically active phenotype identified above. Dome-derived CAFs aligned particularly strongly with pericyte-, mitochondrial- and BMP-associated programs, consistent with the *PDGFRB*- and *NOTCH3*-expressing states enriched in this culture. Lastly, SHOCO CAFs showed the broadest correspondence across the atlas, with prominent *FOS*/*JUN*, matrix/stem-niche and ECM-associated programs alongside *CCL8*, interferon-responsive and inflammatory states, extending the matrix-remodeling and environment-responsive features identified at the cluster level. To identify co-regulated gene modules without relying on discrete cell clustering, we performed unbiased consensus non-negative matrix factorization (cNMF)^56^ on the stromal compartment, which yielded 14 distinct programs (**Figures 2I, 2J** and **S3H**). Among these, SHOCO CAFs preferentially engaged programs NMF2 and NMF4, centered on ECM organization, collagen remodeling, cell–matrix adhesion and tissue interaction, as well as NMF8, dominated by translation and ribosome biogenesis with additional hypoxia- and NF-κB-associated features. Taken together, cluster, pathway, reference-atlas and unbiased program analyses show that SHOCOs shift expanded CAFs beyond the restricted contractile state of 2D culture toward a broader CRC-like stromal repertoire, while enriching ECM-remodeling, hypoxia-responsive and inflammatory programs that distinguish them from conventional dome co-culture.

To identify the upstream drivers of the transcriptional shift in SHOCO versus rBM-Dome cultures, we performed NicheNet analysis^57^ (**Figure S3I**) to assess the target gene regulatory potential of ligands originating from stromal sender cells. This revealed *TGFB1* as the top-ranked prioritized driver. Other highly active stromal ligands included angiogenic factors such as *ANGPTL4* and *ANG*, as well as structural ECM components (*COL3A1*, *COL6A3*, *COL7A1*, *COL13A1*, *COL24A1*, *TNC*, and *EDIL3*).

### SHOCOs maintain tumor-like immune cell numbers, states and interactions

Given their apparently higher structural fidelity to parental tumors compared with PDOs and PDO-CAF co-cultures, we sought to evaluate SHOCOs as immuno-oncology models, which necessitated the inclusion of immune cells. To render SHOCOs immunocompetent, we incorporated peripheral blood mononuclear cells (PBMCs), by including them within the non-polymerized gel droplets, along with PDOs and CAFs (**Figure 1A**). We found that both CD4^+^ and CD8^+^ T cells were retained within SHOCOs over at least five days. At baseline, T cells were typically excluded from the tumor nests, and interspersed within the CAF-rich stromal compartment (**Figure 3A**).

**Figure 3.**
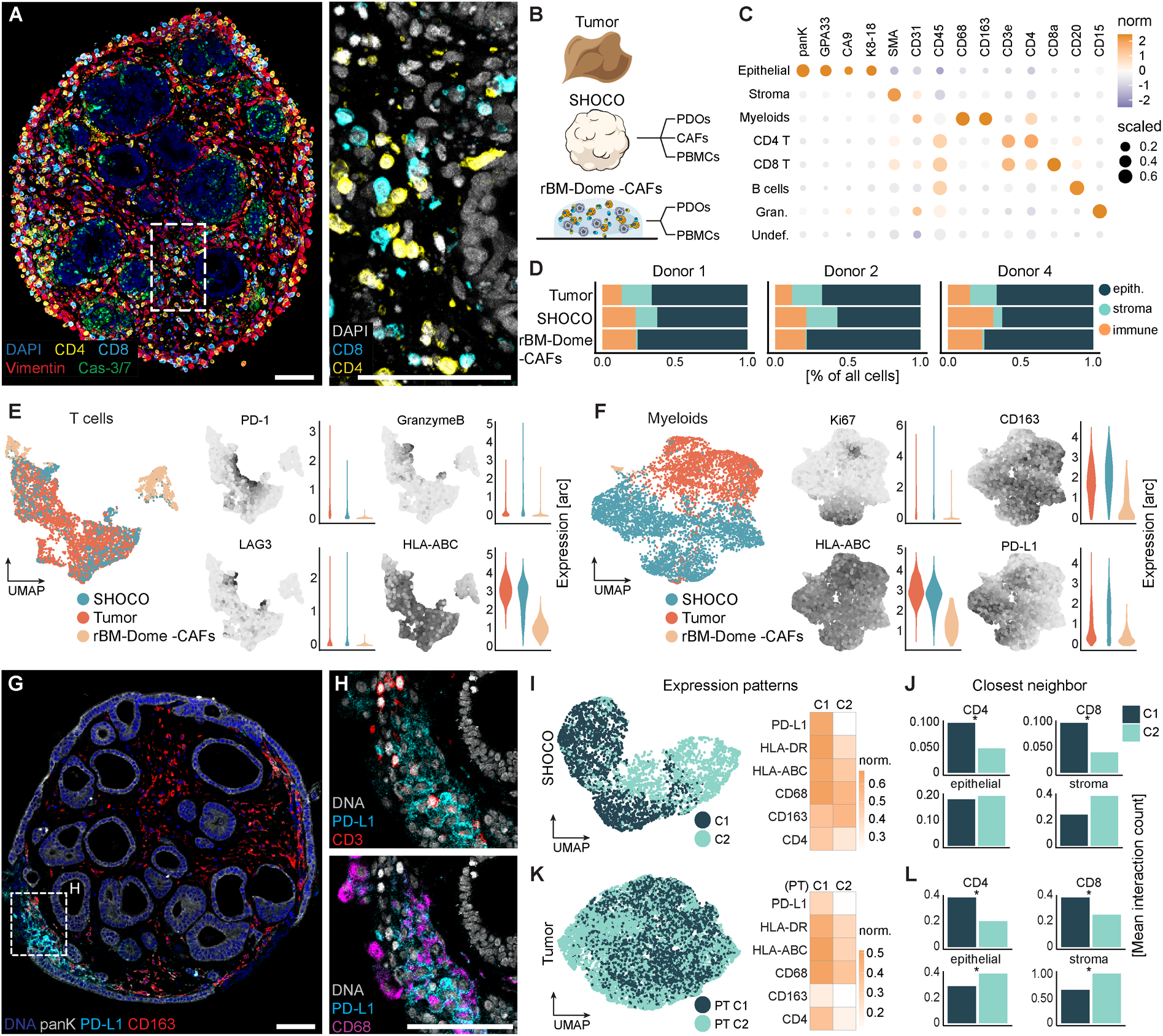
**Compared to generic organoid-PBMC co-cultures, SHOCOs maintain a more faithful immune compartment similar to the primary TME including an extensive myeloid population that mirrors the functionality of its primary counterpart**. **(A)** Multiplex immunofluorescence image of a day 5 SHOCO (tumor Donor 5, allogeneic PBMC donor IC01) showing CD4 and CD8 T cells within the fibroblast-rich stromal compartment, with a high-magnification view of the indicated region. **(B)** Systems compared by spatial mass cytometry: primary tumors, SHOCOs containing matched PDOs and CAFs with allogeneic healthy-donor PBMCs, and rBM-Dome cultures containing PDOs and PBMCs. *N* = 3 tumor donors (Donor 1, Donor 2, Donor 4) and 1 allogeneic PBMC donor (IC02). **(C)** Dot plot of lineage-marker expression used to define epithelial (panK, GPA33, CA9, K8-18), stromal (*a*-SMA), myeloid (CD45, CD163 or CD68), T cell (CD45, CD3, CD4 or CD8), B cell, and granulocyte (CD45, CD15) populations. **(D)** Relative abundance of epithelial, stromal, and immune populations in primary tumors and corresponding SHOCO or rBM-Dome cultures from Donor 1, Donor 2 and Donor 4. **(E)** UMAP of T cells across primary tumors, SHOCOs, and rBM-Dome cultures, with expression maps and distributions of PD-1, granzyme B, LAG3, and HLA-ABC. **(F)** UMAP of myeloid cells across the three systems, with expression maps and distributions of Ki-67, CD163, HLA-ABC, and PD-L1. **(G)** Reconstructed spatial mass cytometry image of a SHOCO (Donor 1) showing the spatial distribution of PD-L1- and CD163-expressing myeloid populations. **(H)** High-magnification views of the region indicated in (G), showing PD-L1 (cyan) with CD3 (red; top) or CD68 (magenta; bottom). **(I)** UMAP analysis of SHOCO myeloid cells identifying an antigen-presenting C1 population enriched for PD-L1, HLA-DR, HLA-ABC, and CD68 and a C2 population characterized by higher CD163 expression. **(J)** Closest-neighbor analysis of C1 and C2 myeloid cells with CD4 T cells, CD8 T cells, epithelial cells, and stromal cells in SHOCOs, shown as mean interaction counts per myeloid cell. **(K)** UMAP analysis of myeloid cells from 52 primary colorectal tumors identifying PT C1 and PT C2 populations and their defining expression patterns. PT C1 is enriched for PD-L1, HLA-DR, HLA-ABC, and CD68, whereas CD163 expression is low in both populations. For (J) and (L), differences between C1 and C2, and PT C1 and PT C2 were assessed separately for each neighboring cell population using permutation tests; the corresponding null distributions and observed differences are shown in Figure S4H. Scale bars, 100 μm.

With immune cells successfully retained within SHOCOs, we next asked whether the model also preserved the phenotypic complexity of the native immune TME. We benchmarked SHOCOs against parental tumors and the widely used PDO-PBMC co-culture models (rBM-Dome(–CAF))^27^. We performed immuno-phenoscaping of the SHOCOs using a highly multiplexed spatial proteomics approach, allowing us to visualize and quantify 40 protein markers at single-cell scale within SHOCOs, parental tumors and rBM-Dome(–CAF) (**Figures 3B** and **S4A–S4B**)^58,59^. The antibody panel was designed to broadly characterize the immune compartment while also distinguishing major epithelial, stromal and endothelial cell populations (**Figure S4C and S4D**). Briefly, we labeled formalin-fixed paraffin-embedded (FFPE) slides containing SHOCOs from three different patient samples, the corresponding primary tumors, and matching conventional co-cultures with metal-tagged antibodies. Following laser ablation, mass spectrometry-based detection and computational reconstruction, we performed single-cell segmentation of the resulting 40-plex-stained images, using commonly accepted markers to attribute cells to an epithelial (panK, GPA33, CAIX, K8-18), stromal (*a*-SMA) or immune lineage (CD45^+^). The marker panel allowed us to further split the immune population into T cells (CD3, CD4 or CD8), B cells (CD20), myeloid cells (CD163 or CD68) and granulocytes (CD15) (**Figure 3C**). The analysis revealed the presence of a robust stromal cell population within the SHOCOs and primary tumors, whereas stromal cells were unsurprisingly absent within the rBM-Dome(–CAF) model (**Figure 3D**).

Comparing the size of the T-cell compartment within the *in vitro* models to that of the primary tumor is of arguably limited significance, considering that we can control the concentration of T cells we include as we construct the models. Instead, we assessed how the emergent T-cell states within the SHOCO model relate to those of their primary counterpart, as compared with traditional rBM-Dome(–CAF). T cells within SHOCOs appeared more phenotypically similar to those within parental samples than T cells within rBM-Dome(–CAF) (**Figure 3E**). This resemblance was underpinned by expression of PD-1 and LAG3, together with granzyme B (GZMB) and HLA-ABC, pointing to a moderately activated state with cytotoxic potential. We examined this phenotype in greater detail by flow cytometry, comparing SHOCOs with compositionally matched collagen-(Col-Dome) and rBM-based dome (rBM-Dome) cultures (**Figure S4E**). Across all systems, CD4^+^ and CD8^+^ T cells progressively acquired activation-associated markers, including CD69, CD25, GZMB and PD-1, with collagen-based cultures generally showing stronger responses than their rBM-based counterparts (**Figure S4F**). FOXP3 remained scarce within the CD4^+^ compartment. One phenotype, however, emerged specifically within SHOCOs: CD8^+^ T cells progressively gained CD103 toward the end of culture, suggesting a shift toward a tissue-residency-associated state known to be promoted by TGF-β signaling in the tumor microenvironment^60,61^.

Aside from T-cell states, we noted unexpected differences in the presence and phenotypic properties of the myeloid compartment between SHOCOs and rBM-Dome(–CAF). In particular, despite incorporating an equivalent fraction of myeloid cells at model construction, SHOCOs were more effective at preserving them, whereas conventional rBM-Dome(–CAF) co-cultures were nearly devoid of myeloid cells by day 5 (**Figure 3F**). Beyond their mere presence, myeloid populations within SHOCOs displayed higher concordance to matching primary samples in the expression of PD-L1, CD163 and HLA-ABC (**Figure 3F**). Flow cytometry further confirmed monocyte-to-macrophage differentiation, with CD64+ cells at day 2 showing high expression of CD80, CD86, CD163 and CD206, consistent with an early macrophage differentiation state rather than distinct polarized populations^22^ (**Figure S4G**).

Within SHOCOs, spatial proteomics uncovered a further layer of myeloid heterogeneity at later time points. Rather than being co-expressed, CD163 and PD-L1 marked distinct populations that appeared spatially segregated (**Figures 3G** and **3H**). A two-cluster separation incorporating all myeloid markers confirmed the presence of two well-differentiated states (**Figure 3I**). The first population (C1), in addition to expressing PD-L1, featured high levels of HLA-ABC and HLA-DR, pointing to a state associated with antigen presentation and immune response^62,63^. The second population (C2) showed lower expression of these markers but was enriched in CD163, consistent with a more macrophage-like phenotype^64–67^.

The functionality of markers involved in antigen presentation and T-cell-mediated immune responses necessitates physical contact between antigen-presenting cells and T cells. To investigate whether the putative antigen-presenting myeloid cell population within the SHOCOs might engage in functional interactions, we performed a nearest-neighbor analysis, mapping the immediate interaction partners of both myeloid clusters (**Figures 3J** and **S4H**). Indeed, we identified a preferential spatial association between both CD4^+^ and CD8^+^ T cells and the antigen-presenting C1 population, compared to C2. The latter, on the other hand, was more likely to associate with stromal cells, whereas both clusters contacted epithelial cells to a comparable extent. To examine the physiological relevance of the myeloid cell states and interactions recorded in SHOCOs, we extended the 40-plex staining and single-cell analysis to an additional 52 primary CRC tumor (PT) samples (**Figure 3K**). Within primary tumors, we likewise identified a myeloid cell population marked by substantial expression of PD-L1, HLA-ABC and HLA-DR (PT C1), and a population expressing lower levels of these markers (PT C2) (**Figure 3K**), which appeared to correspond to C1 and C2, respectively, within SHOCOs (**Figure 3I**). We note that myeloid cells within primary tumors did not form two distinct clusters within the UMAP, which we attribute to the low detection of CD163 across all samples. Remarkably, cellular interaction patterns were likewise conserved (**Figures 3L** and **S4H**): T cells preferentially associated with PT cluster 1, whereas epithelial and stromal cells interacted more extensively with PT cluster 2, in line with observations made in SHOCOs. Altogether, these data highlight key advantages of SHOCOs as a model of the native immune TME, particularly as compared with conventional co-culture models. With these insights in mind, we sought to explore and validate SHOCOs as a model for preclinical immuno-oncology research and drug profiling.

### Immunocompetent SHOCOs elicit robust anti-tumor inflammation in response to cancer immunotherapy drugs

T-cell bispecific antibodies (TCBs) are immunotherapeutic compounds that recognize two different targets: a cancer antigen and the CD3 T-cell receptor (TCR). By crosslinking the target cell and a T cell, TCBs trigger cytotoxic T-cell responses without the need for direct antigen-TCR binding^68–70^. We have previously utilized rBM-Dome(–CAF) co-cultures for joint efficacy-safety profiling of TCBs, showing that the approach captures aspects of clinical outcomes in patients^27^. Compared with conventional co-cultures, SHOCOs combine a stromal compartment with more physiologically representative T- and myeloid cell states (**Figures 3E** and **3F**), while retaining the scale required for drug testing. We therefore set out to evaluate SHOCOs as a platform for the evaluation of TCB anti-tumor effects. We initially modeled an immune-rich, or “hot” TME, expecting substantial TCB-mediated T-cell activation that would allow us to investigate the magnitude, kinetics and characteristics of the anti-tumor response.

We evaluated immune-mediated epithelial damage within SHOCOs through machine learning-mediated segmentation of the histological sections and quantification of the cleaved caspase-3/7. We assessed the efficacy of an EpCAM-targeted TCB (EpCAM-TCB) at different concentrations (0.01, 0.1, 1 µg/mL) and at three different time points (day 1, day 3, day 5) (**Figure 4A**). TCB treatment led to the induction of caspase 3/7 signals within the tumor nests, which increased with time and culminated in complete epithelial destruction by the end of the treatment (**Figures 4B–4J**). The kinetics of TCB-mediated tumor targeting accelerated with higher TCB concentrations. Epithelial destruction was associated with (and typically preceded by) intraepithelial infiltration of CD8^+^ T cells. Conversely, tumor nests within control SHOCOs remained intact, devoid of T cells, and grew and disseminated outward, ultimately coating the edge of the SHOCOs in a manner reminiscent of invasive tumor spread^71,72^. The quantification yielded results consistent with our observations above. Highest TCB concentrations led to rapid induction of apoptosis signals which peaked early (Day 3), followed by a decrease that is likely indicative of complete epithelial destruction (**Figure 4K**). Organoid damage was delayed in the low-concentration condition: the analysis revealed a trend of monotonic caspase-3/7 signal increase, with a peak that is probably reached after Day 5. These data suggest that our model captures not only differences in magnitudes of immune-mediated cytotoxic responses, but also differences in kinetics. Consistent with the decay of epithelial structures observed earlier, the average EpCAM signal also showed a constant downtrend for samples treated with the TCB (**Figure 4K**).

**Figure 4.**
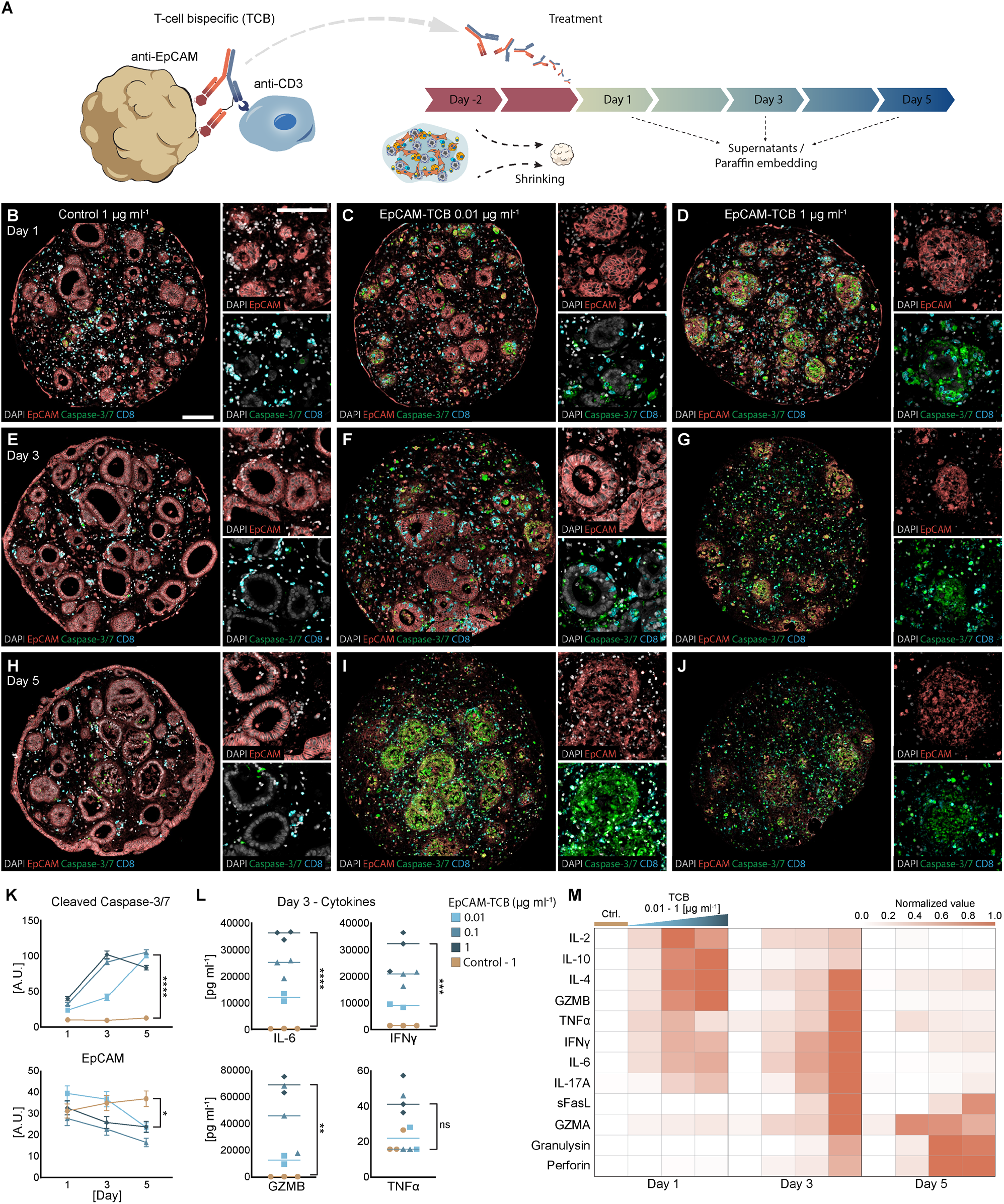
EpCAM-targeted T-cell bispecific treatment induces sustained tumor cell killing and inflammatory immune activation in immunocompetent SHOCOs. **(A)** Schematic of EpCAM-targeted T-cell bispecific (TCB) activity and experimental timeline. SHOCOs were treated after 2 days of culture, with supernatants collected and cultures processed for histological analysis at days 1, 3, and 5 after treatment. **(B–J)** Multiplex immunofluorescence images of SHOCOs treated with control TCB (B, E, H), 0.01 μg/mL EpCAM-TCB (C, F, I), or 1 μg/mL EpCAM-TCB (D, G, J) at days 1 (B–D), 3 (E–G), and 5 (H–J). EpCAM marks epithelial cells (red), cleaved caspase-3/7 apoptotic cells (green), and CD8 T cells (cyan). The control TCB lacks EpCAM specificity and does not productively engage CD3. **(K)** Quantification of cleaved caspase-3/7 and EpCAM signals over 5 days of treatment. SHOCOs from donor 1 were analyzed in triplicate, with five FFPE sections quantified per replicate. Data are shown as mean with SEM. **(L)** Quantification of IL-6, IFNγ, granzyme B, and TNFα in culture supernatants at day 3 across control and EpCAM-TCB treatment concentrations. Data is shown as median with replicates. **(M)** Heatmap of normalized cytokine and cytotoxic effector levels in culture supernatants across EpCAM-TCB concentrations at days 1, 3, and 5. Values were normalized independently for each analyte. Data shown are from tumor Donor 1 (*N* = 1) and allogeneic PBMC donor IC03 (*n* = 3 SHOCOs; 6 FFPE images per SHOCO). Panel (K) was analysed by ordinary one-way ANOVA, and panel (L) by ordinary two-way ANOVA including the interaction between factors, followed by Tukey’s multiple-comparisons test (single pooled variance). Only comparisons between EpCAM-TCB (1 µg/mL) and the corresponding control are annotated. P ≥ 0.05 (ns); P < 0.05 (*); P < 0.01 (**); P < 0.001 (***); P < 0.0001 (****). Scale bars, 100 μm.

In parallel with quantifying organoid apoptosis, we evaluated TCB-mediated T-cell activation by measuring the secretion of common pro-inflammatory cytokines at the three experimental time points. We employed a 13-plex assay using beads to capture free floating cytokines from the supernatant, which were then analyzed by flow cytometry. We found that key cytokines associated with inflammation (IL-6, IFNγ) and cytotoxicity (GZMB) were induced at high levels and in a TCB-concentration manner (**Figure 4L**). We further explored the kinetics of cytokine production, and observed an early (Day 1 of TCB treatment) appearance of GZMB, IL-2 and IL-4, whereas cytokines including GZMA, IFNγ, IL-6 and IL-17A peaked at the intermediate time (Day 3) post treatment. Finally, the analysis revealed late (Day 5) induction of perforin and granulysin (**Figure 4M**). Thus, we demonstrate that the SHOCO model is amenable to quantitative and longitudinal analyses of parameters commonly used preclinically as readouts and clinically as biomarkers of anti-tumor responses^70,73,74^.

We next benchmarked SHOCOs as a platform for TCB profiling against other PDO-based culture formats, focusing on the magnitude and kinetics of the TCB response (**Figure 5A**). Cytokine profiling of SHOCOs, collagen- (Col-Dome) and rBM-based (rBM-Dome) PDO-CAF-PBMC domes, and conventional PDO-PBMC co-cultures (rBM-Dome(–CAF)) revealed prominent differences in immune activation across both EpCAM- and CEACAM5-targeting TCBs. SHOCOs consistently produced the strongest cytokine response, followed by collagen-based domes, whereas both rBM-based systems showed a more modest response (**Figures 5B** and **5C**). Notably, IL-6 was virtually absent from the rBM-Dome(–CAF) co-culture, consistent with an important contribution of both fibroblasts and myeloid cells to the inflammatory response^75,76^.

**Figure 5.**
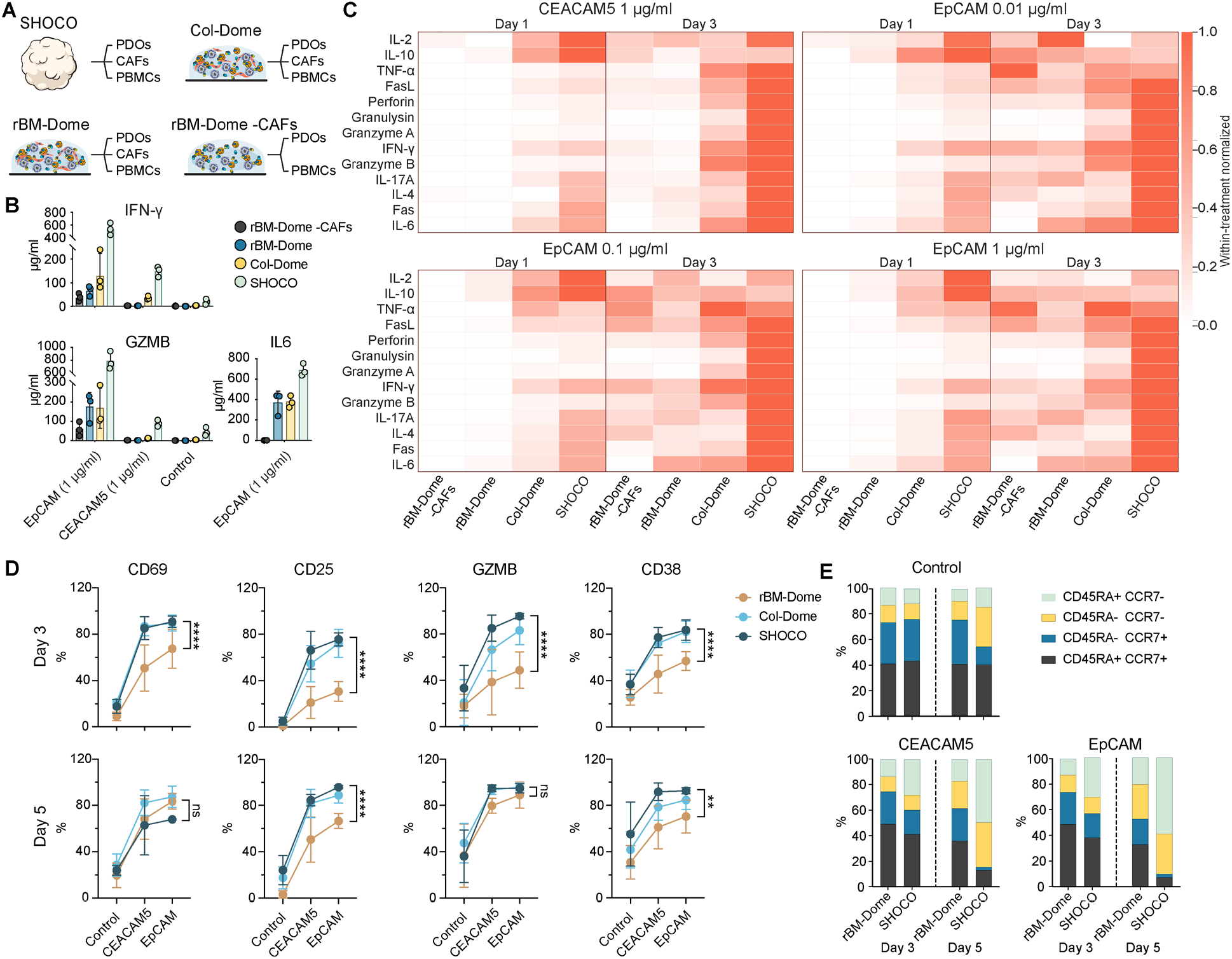
SHOCOs mount rapid and sustained responses to T-cell bispecific treatment, with enhanced cytokine release, T-cell activation, and effector differentiation compared with conventional dome cultures. (A) *In vitro* systems compared following T-cell bispecific (TCB) treatment: SHOCOs, collagen-and rBM-based dome cultures containing PDOs, CAFs, and PBMCs, and conventional organoid-PBMC co-cultures lacking CAFs. **(B)** Raw cytokine concentrations in supernatants at day 3 for Donor 5 across all four systems. IFN-γ and granzyme B (GZMB) are shown following EpCAM- and CEACAM5-targeting treatments (1 µg/mL) or control treatment, whereas IL-6 is shown only for the EpCAM-targeting treatment. Circles represent individual replicate wells and bars show mean ± SD. **(C)** Heatmaps of cytokine and cytotoxic effector secretion across the four culture systems at days 1 and 3 following treatment with CEACAM5-TCB (1 μg/mL) or EpCAM-TCB (0.01, 0.1, or 1 μg/mL). Values were normalized within each treatment condition. *N* = 2 tumor donors (Donor 1, Donor 5) and 1 allogeneic PBMC donor (IC04); *n* = 3 cultures per donor per system. **(D)** Flow cytometry assessment of T cell activation markers CD69, CD25, granzyme B, and CD38 in rBM-based dome cultures, collagen-based dome cultures, and SHOCOs at days 3 and 5 following control, CEACAM5-TCB (1 µg/mL), or EpCAM-TCB (1 µg/mL) treatment. Data is shown as mean ± SD and analysed by ordinary two-way ANOVA including the interaction between factors, followed by Tukey’s multiple-comparisons test (single pooled variance). Comparisons between SHOCO and rBM-Dome cultures are annotated. **(E)** Distribution of CD8 T-cell differentiation states defined by CD45RA and CCR7 expression in rBM-based dome cultures and SHOCOs at days 3 and 5 following control, CEACAM5-TCB (1 µg/mL), or EpCAM-TCB (1 µg/mL) treatment. P ≥ 0.05 (ns); P < 0.05 (*); P < 0.01 (**); P < 0.001 (***); P < 0.0001 (****).

Flow cytometry further revealed that these differences were driven in part by response kinetics: by day 3, CD8^+^ T cells in both collagen-based systems already displayed strong induction of CD69, CD25, GZMB and CD38, while activation in rBM-based cultures developed more gradually and largely caught up by day 5 (**Figure 5D**). TCB stimulation also reshaped the CD8^+^ T-cell compartment within SHOCOs, increasing the proportion of CCR7^−^ cells and, in particular, CD45RA^−^CCR7^−^ and CD45RA^+^CCR7^−^ populations (**Figure 5E**). The latter phenotype is commonly associated with terminally differentiated effector-memory cells, although its emergence here may simply reflect the prolonged T-cell stimulation imposed by continuous TCB-mediated antigen-CD3 crosslinking. Intriguingly, this shift coincided with the late appearance of a strongly cytotoxic effector profile, dominated by perforin and granulysin and accompanied by GZMA. Coordinated expression of these cytotoxic molecules has been associated with advanced effector-memory differentiation in human CD8^+^ T cells^77,78^, further suggesting that prolonged TCB stimulation drives the cells toward a highly differentiated cytotoxic state. Together, these comparisons indicate that while T-cell activation can be elicited across all culture formats, collagen-based environments accelerate its onset, with the tissue-like SHOCO setting supporting the strongest overall immune response. Remarkably, the dense, compact architecture of SHOCOs contributed to elevated susceptibility to T-cell activation beyond effects underpinned by CAF presence *per se*: SHOCOs appeared more immunogenic than CAF-containing, collagen-based dome cultures, despite identical cellular and ECM compositions at construction.

### A window into the tumor stroma: CAFs as modulators of T-cell behaviors and immunotherapy targets

The dense fibrotic stroma of solid tumors has long been implicated in undercutting anti-tumor immunity by limiting immune-cell access to the tumor epithelium or via diverse other immunosuppression mechanisms^79–85^. Surprisingly, CAF presence within conventional co-culture did not attenuate but instead appeared to enhance TCB responses (**Figures 5A** and **5B**), possibly because mere inclusion of fibroblasts in loose dome culture does not fully recapitulate their physical and steric effects, while capturing their inflammatory functions. We hypothesized that the dense stromal structure of SHOCOs would be more effective in capturing CAF-mediated influences on anti-tumor immunity. We tuned the stromal compartment of SHOCOs to examine how stromal abundance influences TCB-mediated anti-tumor responses. SHOCOs containing either 5000 or 25000 CAFs per microliter were treated with a CEACAM5-targeting TCB, whose lower potency compared with EpCAM-TCB afforded a longer window in which to follow immune-cell infiltration and tumor killing^27,86^. As expected, treatment induced caspase-3/7 within epithelial niches and progressive accumulation of CD8^+^ T cells (**Figures 6A** and **6B**). Quantification of cleaved caspase-3/7 revealed no significant difference between low- and high-stroma conditions, indicating that cytotoxic activity was preserved despite increased stromal density (**Figure 6C**). This was reflected in the epithelial compartment, where TCB treatment further reduced the already smaller niches in CAF-high SHOCOs, resulting in the lowest epithelial area overall (**Figure 6D**). However, despite a comparable acute cytotoxic response, we found that stroma density influenced T-cell behaviors that are likely to shape longer-term responses. By day 3 of treatment, CAF-high SHOCOs harbored significantly lower numbers of intraepithelial T cells, compared with CAF-low SHOCOs, suggesting that high stromal density impedes intraepithelial T-cell infiltration (**Figure 6E**). To assess the relevance of these findings to real tumors, we queried the relationship between intraepithelial T-cell presence and density of the adjacent stroma within our cohort of 52 primary CRC samples profiled by spatial proteomics. We found that increasing stromal abundance correlated with fewer T-cell-epithelial interactions, closely mirroring the exclusion phenotype observed in CAF-high SHOCOs (**Figure 6F**). Flow cytometry further showed that this reduced access translated into weaker T-cell activation by day 5: CD69, CD25 and GZMB were all significantly higher in T cells from low-stroma SHOCOs than in their CAF-high counterparts (**Figure 6G**).

**Figure 6.**
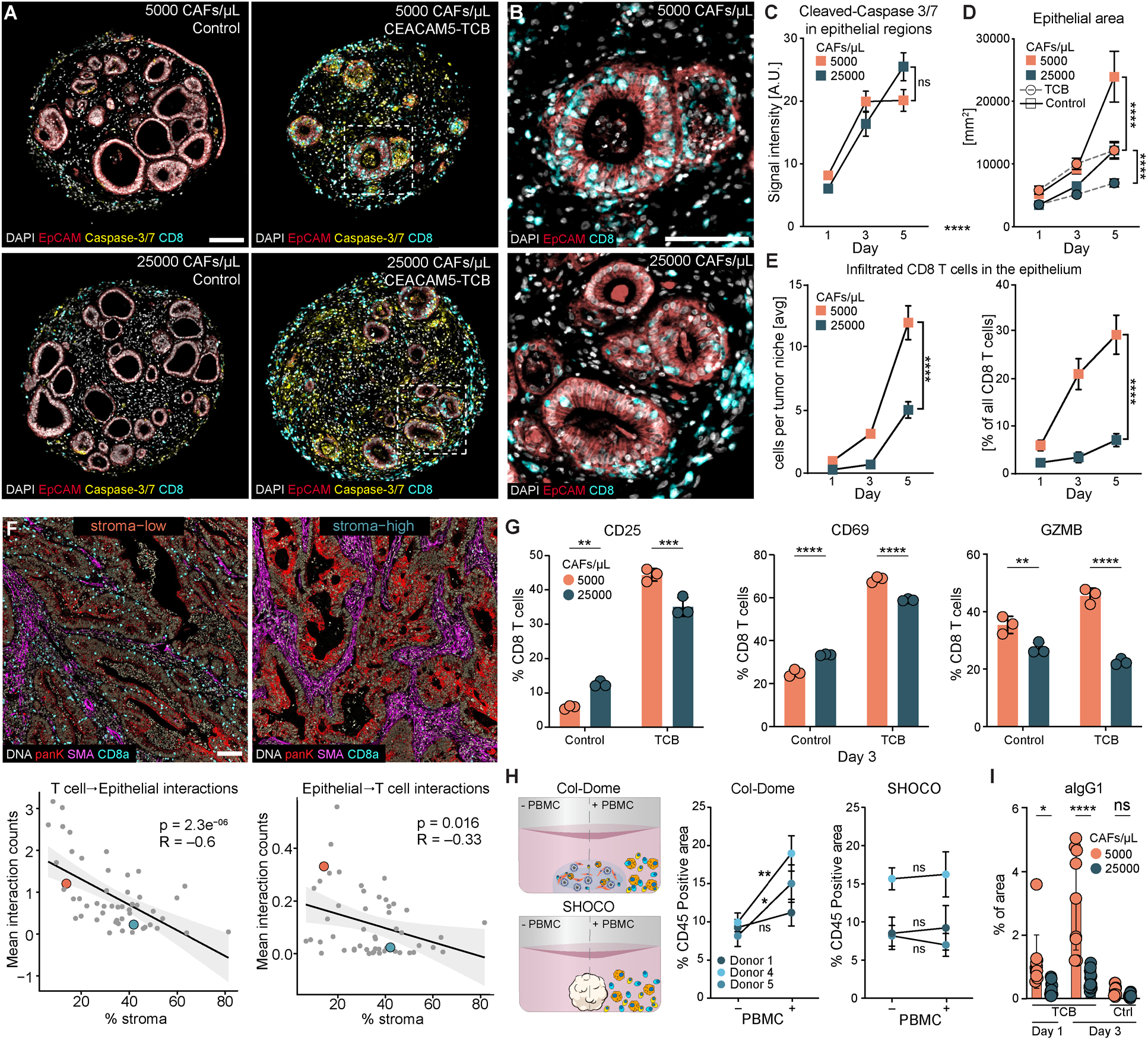
SHOCOs reveal stromal compaction as a physical determinant of T-cell and TCB access, recapitulating immune exclusion observed in primary CRC. **(A)** mIF images of SHOCOs (tumor Donor 1 and allogeneic PBMC donor IC05) generated at low (5000 CAFs/µL) or high (25000 CAFs/µL) stromal density after 5 days of CEACAM5-TCB treatment or Control. **(B)** High-magnification images of the CEACAM5-TCB-treated SHOCOs from panel (A) showing CD8 T-cell infiltration into epithelial niches at low and high stromal density. **(C)** Quantification of cleaved caspase-3/7 signals over 5 days of treatment in SHOCOs generated at low or high stromal density. **(D)** Quantification of epithelial niche area over 5 days in control- and CEACAM5-TCB-treated SHOCOs at low or high stromal density. **(E)** Quantification of CD8 T-cell infiltration into epithelial niches over 5 days, shown as total infiltrating CD8 T cells and as the fraction of all CD8 T cells within SHOCOs. For (C-E), data were pooled from four independent experiments comprising three tumor donors (Donor 1, tested twice; Donor 3; Donor 7) and two allogeneic PBMC donors (IC05 and IC06). Each experiment included three SHOCOs per condition, quantified from approximately five immunostained FFPE sections per SHOCO. Data shown as mean with SEM and analysed by ordinary two-way ANOVA including the interaction between factors, followed by uncorrected Fisher’s LSD tests (single pooled variance). **(F)** Reconstructed spatial mass cytometry images of representative primary colorectal tumors with low (red) or high (blue) stromal *a*-SMA density. Regression analyses include all 52 tumors from this independent cohort and show the relationship between stromal *a*-SMA-positive area and mean T-cell-epithelial or epithelial-T-cell interaction counts. **(G)** Flow cytometry quantification of CD25, CD69, and granzyme B expression in CD8 T cells at day 3 following CEACAM5-TCB treatment or Control in SHOCOs generated at low or high stromal density. *N* = 1 tumor donor (Donor 1) and 1 allogeneic PBMC donor (IC07); *n* = 3 SHOCOs per condition. **(H)** Experimental design comparing compositionally matched Col-Dome cultures and SHOCOs generated with or without externally added PBMCs. Quantification of % CD45-positive area after 3 days of EpCAM-TCB (0.1 µg/mL) treatment. *N* = 3 tumor donors (Donor 1, Donor 4, Donor 5) and 1 allogeneic PBMC donor (IC07); *n* = 3 per condition, mean ± SD. Statistical significance was evaluated using Welch’s two-tailed independent t-test. **(I)** Quantification of anti-IgG1-positive area in control- and TCB-treated SHOCOs at days 1 and 3, comparing low and high stromal density. *N* = 1 tumor donor (Donor 1) and 1 allogeneic PBMC donor (IC07); *n* = 10-12 SHOCOs per condition; 2 FFPE images per SHOCO. For (G) and (I) data were analysed by ordinary two-way ANOVA including the interaction between factors, followed by Šídák’s multiple-comparisons test (single pooled variance). P ≥ 0.05 (ns); P < 0.05 (*); P < 0.01 (**); P < 0.001 (***); P < 0.0001 (****). Scale bars, 100 μm.

The mechanisms by which the stroma shapes anti-tumor immune responses are complex and manifold^85,87,88^. The most widely established modes of stroma-mediated immunosuppression are physical effects, whereby stromal cells and ECM physically impede immune cell infiltration, or signaling effects, whereby stromal factors, such as TGF-β, actively suppress immune cell functions^83,84^. Having established a system that captures stromal influences on immunotherapy responses, we set out to elucidate the mechanisms behind the attenuated inflammatory response within the high-density SHOCOs. We first focused on the physical and mechanical role of the stroma. Starting from identical cellular and ECM compositions, we formed floating SHOCOs or dome structures attached to the tissue culture substratum, which, in the latter, prevents CAF-mediated compaction into a dense and mechanically stiff tissue (**Figure 6H**). To examine the effect of this stromal physical “barrier” occurring in the floating SHOCOs on T-cell recruitment, we generated an immune-poor, or “cold”, TME in which only a small fraction of PBMCs was incorporated into the tissue to create an initial inflammatory trigger driving subsequent immune recruitment. The remaining immune cells were introduced into the surrounding medium and their influx into the SHOCO was monitored and quantified. We observed a pronounced reduction of infiltrated immune cells within compact SHOCOs, compared with their sparse counterparts (**Figures 6H** and **S5A**), despite an identical cellular and ECM composition at construction. This difference suggests that mechanical hindrance is a key mechanism by which the stroma influences immune cell infiltration in the system. In line with this conclusion, we observed lowest immune-cell infiltration in Col-Dome cultures constructed from the PDO-CAF donor that displayed the strongest contractile capacity (**Figure S2D**), which further underscores the critical role of tissue compaction and mechanics in stroma-mediated immune exclusion.

Given the established immunoregulatory role of TGF-β, we additionally considered TGF-β-mediated immunosuppression as another mechanism through which the stroma influences T-cell activity in the system. We first assessed TGF-β presence and stromal dependence, detecting robust levels of extracellular TGF-β, which increased further in CAF-high SHOCOs (**Figure S5B**). To evaluate whether stromal suppression of T-cell activation could be rescued by TGF-β inhibition (**Figure S5C**), we combined TCB treatment with the TGF-β-neutralizing antibody fresolimumab (**Figure S5D**) or the TGF-β receptor inhibitors galunisertib and SB431542. Aside from subtle effects early upon TCB treatment, TGF-β inhibition failed to durably rescue the suppression of T-cell activation occurring within CAF-high SHOCOs. Thus, despite increased production, TGF-β does not appear to be a major mechanism of stroma-mediated immunosuppression within the model.

Beyond restricting immune-cell movement, dense stromal architecture has been shown to limit therapeutic antibody penetration^54^, which would ultimately lead to diminished T-cell-target engagement and inflammatory activation. In primary tumors, stroma-mediated hindrance of antibody access was linked to the abundance of periostin^54^, which was transcriptionally upregulated specifically in SHOCOs (**Figure S3D**). To consider this mechanism, we assessed TCB tissue distribution and target binding within CAF-low and -high SHOCOs. CEACAM5-TCB signal increased between days 1 and 3 but remained significantly lower in CAF-high SHOCOs at both time points, confirming that stromal density impedes TCB penetration (**Figures 6I** and **S5E**). We note that the distribution of the signal was consistent with the pattern of CEA expression in tumors and PDOs^27^, and its intensity was higher in peripheral epithelial niches (**Figure S5E**). The control TCB produced no detectable staining, which is consistent with the absence of a target-binding arm required for TME retention. These data suggest that elevated stromal density dampens anti-tumor TCB action through simultaneous physical hindrance of both T-cell migration and therapeutic antibody penetration, mirroring observations in patient tumors.

As shown above and suggested across the literature, cancer-associated fibroblasts may undercut the effectiveness of endogenous anti-cancer immunity and cancer immunotherapy. Given their abundance in tumors, novel therapeutic approaches aim to turn CAFs from an obstacle into an advantage, by using them to localize and enhance the effects of T-cell-targeted therapies. Notably, these therapies are challenging to evaluate using tumoroid-immune cell co-cultures, which lack stromal cells, or *in vivo* models, which do not afford the control and modularity required to optimize drug pharmacological properties and lack human-specific isoforms of therapeutic targets. We decided to explore SHOCOs as platforms for the evaluation of CAF-targeted therapies. In particular, we treated SHOCOs with cibisatamab (CEA-TCB^26,70^) alongside the fibroblast-activation protein (FAP)-targeting compound RO7122290 FAP-4-1BBL (**Figure S5F**)^89–91^. FAP-4-1BBL is a bispecific fusion protein carrying a split trimeric 4-1BB (CD137) ligand and a fibroblast-activation protein α (FAP) binding site. This design co-stimulates T cells, enhancing tumor cell killing in FAP-expressing tumors. The combination treatment specifically targets CEACAM5 and FAP-expressing tissues, enhancing the antitumor effects of cibisatamab without raising its concentration or risk of side effects^92^. Cytokine analysis of double-treated SHOCOs revealed elevated levels of key pro-inflammatory cytokines – GZMB, IFNγ, and IL-6 – following CEA-TCB treatment. The addition of FAP-4-1BBL further amplified cytokine secretion, doubling IFNγ concentrations by day 3 (**Figure S5F**) compared to cibisatamab monotherapy as reported in the context of the equivalent clinical trial in peripheral blood of patients^92^. Together, these experiments illustrate how the same stromal compartment that can shield tumors from immune attack may also be turned into a therapeutic advantage, and establish SHOCOs as a versatile platform for studying this interplay in a controlled human TME.

## Discussion

Tumor behavior and response to therapy are shaped not only by malignant cells, but by the cellular, structural and biochemical environment in which they reside^93,94^. Reproducing this complexity in human preclinical models remains challenging, particularly without sacrificing the reproducibility, scalability or experimental control required for mechanistic studies and drug development. We developed shaking organoid co-cultures (SHOCOs), a scalable 3D model that combines patient-derived colorectal tumor organoids (PDOs) with cancer-associated fibroblasts (CAFs), tumor-relevant extracellular matrix (ECM) and immune cells in a mechanically unconstrained culture.

In this model, CAFs are not introduced as simply another cellular component, but are allowed to actively construct their environment. In an untethered type I collagen matrix, CAF-mediated contraction and ECM remodeling reorganize the culture into a dense, heterogeneous structure. This self-organization produces stromal regionalization, hypoxia and a necrotic-like core. Although conventional PDO-CAF (rBM-Dome) co-cultures remain highly useful, our comparisons show that both matrix composition and mechanical freedom strongly influence the biology that emerges across both the stromal and immune compartments. While the starting CAF cultures were dominated by myofibroblastic and proliferative programs, CAFs within SHOCOs became reprogrammed into diverse tumor-relevant subsets that spanned a broader range of stromal programs identified in primary CRC^31^.

Aside from stromal cells, the SHOCO microenvironment shaped the immune cells introduced into the system, influencing their representation, phenotypic and functional states. Whereas myeloid cells introduced into conventional PDO-PBMC co-cultures become rapidly depleted, SHOCOs maintained a robust myeloid compartment, likely instructed by the soluble and biophysical milieu of the system^22,95^. Monocytes acquired a macrophage phenotype and diversified into distinct myeloid states, including an antigen-presentation-associated population that preferentially interacted with T cells. These states and interactions mirrored those found in primary CRC samples, suggesting that SHOCOs provide a tumor-relevant platform in which they can be experimentally interrogated.

T cells within SHOCOs likewise inhabit a distinct baseline state, associated with elevated activation and expression of the tissue-residence marker CD103, which permits engagement with epithelial cells^25,96^. The induction of CD103 suggests that blood-derived T cells begin to acquire features of tissue adaptation within SHOCOs, which may be orchestrated in part by elevated TGF-β signaling^96^ and the distinct tissue-like physical features^97,98^ of the model. This heightened baseline activation state is not wholly unexpected in an allogeneic system and may be reinforced by TNFα-NF-κB signaling and antigen-presentation-associated myeloid cells within SHOCOs specifically. However, baseline activation alone is unlikely to explain the stronger TCB response in SHOCOs. Their compact architecture likely also shapes the latter by increasing cellular proximity and altering the local accumulation and diffusion of soluble mediators compared with the more dispersed dome cultures^99,100^. Autologous PBMCs or TILs would help distinguish alloreactivity from TME-driven immune adaptation and allow tumor-specific immunity to be modeled more directly.

CAF-mediated immunosuppression in CRC encompasses both signaling-dependent mechanisms, including TGF-β and chemokine-mediated immune regulation, and structural mechanisms arising from matrix remodeling and tissue organization^84,101,102^. These processes frequently coexist in primary tumors, making their individual contributions difficult to resolve. Exploiting the system’s modularity and control, we varied CAF density to generate defined stroma-poor and stroma-rich environments, bracketing the stromal densities measured in primary CRC. We show that denser stroma attenuated the TCB response, with reduced T-cell access to epithelial niches and weaker activation. Having captured the suppressive effect of dense stroma on therapy response in an experimentally tractable system, we were able to investigate the underlying mechanisms. Several observations point to a substantial physical component underlying this exclusion. In particular, mechanical tissue compaction was required to impede immune cell infiltration: T-cell infiltration was significantly higher in compositionally matched co-cultures that were prevented from compaction. Furthermore, dense stroma constrained epithelial niche expansion even in the absence of immune cells, pointing to a strong compressive mechanical effect. Finally, an increase in density resulted in reduced TCB penetration into the tissue, indicating that the effects of stromal density extend to the physical accessibility of the tumor by molecular therapeutics^54,103^.

Collectively, these data emphasize the importance of considering tissue and tumor architecture within preclinical drug development models. Previous CRC organoid-CAF co-cultures found that increasing fibroblast numbers did not measurably impair TCB-mediated cytotoxicity^26^, consistent with our observation that CAF presence alone is insufficient to produce immune exclusion. The compact architecture of SHOCOs formed through CAF mechanical activity uniquely enabled the emergence of patient-relevant mechanisms whereby dense stroma undercuts therapy effects. Importantly, both of these mechanisms occur in real tumors, as we and others have shown^54^ . Given that stromal barriers and their effects on therapy are represented in SHOCOs, the model could be used to perform genetic and pharmacological screens aimed at better understanding and overcoming them. Indeed, we provide an example of how tumor stroma can be turned from an impediment into a therapeutic strategy, showing that the fibroblast-targeted costimulatory molecule FAP-4-1BBL enhances TCB-mediated anti-tumor activity. This therapeutic paradigm has recently been validated in the clinic^92^.

Together, these findings establish SHOCOs as a controllable human CRC model that captures tumor-relevant organization, cell states, interactions and therapy responses, while affording the scale and reproducibility required in a preclinical drug development assay. As such, they significantly expand our toolkit for both studies in fundamental CRC biology and translational efforts aimed to treat a disease with urgent unmet medical need.

## Limitations of this study

There are several limitations to this study that should be acknowledged. First, the present implementation of immunocompetent SHOCOs relies primarily on allogeneic PBMCs rather than patient-matched autologous PBMCs or TILs. While allogeneic T cells are fully functional for evaluating TCBs, which engage CD3 independently of native TCR-MHC specificity, the absence of matched tumor antigen recognition limits the direct evaluation of therapies relying on endogenous antigen presentation, such as checkpoint inhibitors or neoantigen vaccines. Pilot attempts to incorporate cryopreserved autologous TILs yielded insufficient viable cells for sufficient replicates of SHOCOs. Consequently, ongoing work is required to refine fresh tissue acquisition strategies and optimize *ex vivo* TIL expansion protocols. Second, a conceptual strength and inherent boundary of the platform is that SHOCOs are designed to construct modular hypothesis driven tumor archetypes simulating controlled variations in stroma density and immune infiltration rather than serve as static patient specific histological replicas. As an *ex vivo* reductionist model, SHOCOs do not capture the systemic physiological complexity of naive colorectal cancer tumors due to the absence of functional vasculature, lymphatics and tertiary lymphoid structures. Finally, the tissue-like density and compaction that define SHOCOs introduce technical trade-offs: the opacity of dense collagen prevents live whole-mount imaging, needing section-based multiplex histopathology or spatial proteomics, while extensive enzymatic digestion required for flow cytometry single-cell dissociation can cleave sensitive surface receptors. Nevertheless, these technical constraints are compensated by the platform’s ability to preserve spatial architecture, maintain intact cell-cell cross-talk, and support drug testing.

## Acknowledgements

We thank the patients who generously donated tissue samples used in this study. We thank Salvatore Piscoglio for overseeing the acquisition of primary colorectal tumor samples and associated clinical information from the University Center for Gastrointestinal and Liver Disease (Clarunis), Basel, and Humanitas Research Hospital, Milan, and for the establishment of patient-derived organoid lines and donor-matched cancer-associated fibroblasts. We thank Andrea Corsinotti and colleagues at the 360 Labs Genomics Platform (pRED, Roche) for their support with the single-cell sequencing experiments, and the Flow Cytometry team for their support with cell sorting and sample preparation prior to single-cell library generation. We thank Tamara Tanos for her guidance and feedback on experiments involving FAP-4-1BBL and cibisatamab. We also thank Emma Vidal (DrawImpacts) for the illustration of the graphical abstract. We thank Nadine Stockar and José Galvan for providing valuable guidance and experimental protocols for the analysis of TCB distribution within histological sections and Hartmut Koeppen for comparing SHOCOs and their parental tumor histological sections.

## Author contributions

A.M.F., N.G., and L.C. conceived and designed the study. A.M.F. performed the majority of the experiments and analyses. B.L. and A.H. performed experiments and contributed to data analysis. K.K., M.F.H., I.L., and W.J. contributed to experimental work and supported the study. I.C. led all histology-based work, including Histogel and paraffin embedding, FFPE sectioning, multiplex immunofluorescence staining, and slide scanning. S.T. assisted with Histogel embedding, FFPE sectioning, IHC staining, and slide scanning. A.M.F. analyzed mIF images using HALO AI under the guidance of M.F.H., and I.L. assisted with microscopy and mIF analysis. P.J.-M. and F.C. optimized the hypoxia-assessment protocol. E.D. and M.D. performed staining and acquisition for imaging mass cytometry, while B.E. and S.C. analyzed the spatial proteomics data. F.C. and S.Co. collaborated throughout the study. M.C.-L. coordinated the sourcing of primary colorectal tumor samples and associated patient information used to establish PDO and CAF cultures.

A.M.F. and B.L. performed the single-cell RNA-sequencing experiments, with S.J. performing 10x Genomics library preparation and sequencing. G.P. and S.M. performed the primary computational analysis of the scRNA-seq data, with A.M.F. and B.L. contributing to downstream analysis, interpretation, and visualization. N.G. and L.C. supervised the study. A.M.F., B.L., N.G., and L.C. wrote the manuscript, with input and approval from all authors.

## Declaration of generative AI and AI-assisted technologies in the writing process

During the preparation of this work, the authors used DeepNetV2 from HALO AI (Indica Labs) for image segmentation and cell phenotyping. Generative AI tools were used to assist with language editing and refinement, literature searches and reference identification, and the processing and organization of selected mIF and cytokine datasets. All AI-assisted outputs were reviewed and verified by the authors, who take full responsibility for the content of the manuscript.

## Code availability

No original code was developed for this study.

## Data availability

The scRNA-seq have been deposited in the Gene Expression Omnibus (GEO) database under the accession numbers: GSE347204 (Access token: arirwemqfvedxyp).

## Methods

### Tumor organoid and CAF mono-culturing

List of donors is specified in Table S2. Human tissues and associated clinical information were obtained from patients undergoing tumor resection at the University Center for Gastrointestinal and Liver Disease (Clarunis), Basel, Switzerland. Patient-derived organoids (PDOs) and cancer-associated fibroblasts (CAFs) were sourced through a collaborative framework adhering to legal and ethical regulations. The project was performed in accordance with the Helsinki Declaration and reviewed and approved by the Ethics Committee of Basel, EKBB, no. 2019-02118. Additional tissues, for the generation of PDOs and CAFs models and associated clinical information were obtained from patients undergoing tumor resection at the Humanitas Research Hospital, Milan, Italy. Samples were sourced through a collaborative framework adhering to legal and ethical regulations. The project was reviewed and approved by Comitato Etico Territoriale Lombardia 5 (Ethical approval 3631).

PDOs were cultured in 24-well plates, each well containing a 25 µL Cultrex dome (3432-005-01, R&D Systems). They were passaged approximately every seven days. Media, consisting of a 1:1 mixture of Advanced DMEM-F12 (12634-010, Gibco) + GlutaMax-I (35050-038, Gibco) + Penicilin/Streptomycin (P/S) (15140-122, Gibco) and unsupplemented OGM (100-0190 StemCell), was exchanged every two days, except over the weekend when a three-day interval was permitted. For passaging, media was removed, and wells were washed with 500 µL of Dulbecco’s Phosphate Buffered Saline (DPBS) (14190-094, Gibco) and 500 µL of Gentle Cell Dissociation Reagent (100- 0485, StemCell) was added to each dome. After a 2-minute incubation at room temperature, domes were disrupted and combined (per patient) into a 15 mL Falcon tube. PDOs were further incubated for 10 minutes at room temperature, with resuspension every 3 minutes to mechanically break down the gel and PDO structures. Samples were centrifuged at 250 g for 4 minutes to remove the supernatant, after which the cells were washed with 10 mL of ice-cold wash buffer, consisting of Advanced DMEM-F12 + 1% BSA (130-091-376, Miltenyi Biotec) + HEPES (15630-056, Gibco) + 1x P/S. If undigested gel remained, the washing step was repeated. Once the cells were free of hydrogel and pelleted, any remaining wash buffer was carefully removed, and cells were resuspended in fresh Cultrex. PDOs were then replated in 25 µL domes, the plate was inverted to prevent sedimentation, and incubated at 37°C for 20 minutes to allow polymerization. After polymerization, 500 µL of Media supplemented with 1 µM Y-27632 Rock-Inhibitor (72305, StemCell) was added to each well.

Cancer-associated fibroblasts (CAFs) were cultured in Falcon Culturing Flasks, either 75 cm² (353136, Falcon) or 175 cm² (353112, Falcon), and passaged once confluence reached 95%. Passaging was performed by removing the old CAF media (C28009, PromoCell), washing once with 10 mL DPBS by carefully adding it to the flask and tilting the flask to rinse the entire cell monolayer. The DPBS was then removed, and TrypLE Express (12604-013, Gibco) was added to the flask, 3 mL for the 75 cm² flask and 5 mL for the 175 cm² flask. The cells were incubated for 6 minutes in an incubator at 37°C. The now-floating CAFs were recovered by adding 10 mL of CAF media and washing the bottom of the flask multiple times, pooling all cells in suspension, and transferring them to a 15 mL falcon tube. Cells were centrifuged at 300 g for 3 minutes, and the supernatant was removed, after which the cells were washed once with 10 mL of CAF media to remove any TrypLE remnants. After the wash, the cells were centrifuged again, resuspended in 10 mL of CAF media, and seeded in a new flask at the desired ratio (e.g., for a 1:5 split, only 2 mL were passaged to the new flask). CAF media was then added to a total volume of 15 mL for the 75 cm² flask, and 30 mL for the 175 cm² flask. If CAFs were needed for experimentation, they were counted from the remaining cell suspension and used accordingly.

Both cell populations were maintained in a standard incubator at 37°C, 21% O₂, and 5% CO₂.

### PBMC freezing & thawing

Healthy donor PBMCs were extracted from buffy coats using SepMate™-50 gradient separation, following the manufacturer’s instructions (85450, StemCell). In a nutshell, the SepMate 50 mL tube’s lower compartment was filled with 15 mL Ficoll-Paque (17144003, Cytiva) followed by 15 mL of blood slowly added into the upper compartment so as not to mix the two. The tube was then filled with DPBS supplemented with 2% heat-inactivated fetal bovine serum (FBS) (A31605-01, Gibco). Tubes were then centrifuged, leading to the target cells remaining in the upper compartment while the lower compartment contained the waste. PBMCs were then further washed, counted, and processed as needed. In the majority of cases the cells were aliquoted at different concentrations and cryopreserved using Cellbanker 1 (11910, Amsbio), then stored in liquid nitrogen.

Thawing of frozen PBMCs was performed by placing the vial in a water bath for approximately 1 minute, until only a small ice sphere remained. The vial was then removed from the water bath, and the contents were resuspended slowly to homogenize the solution. The cells were then transferred, drop-by-drop, to a 15 mL Falcon tube prepared with 10 mL of washing media (Advanced DMEM-F12 + P/S + HEPES + 1% BSA). The cells were centrifuged at 500 g for 3 minutes, the supernatant was removed, and the cells were washed once with 10 mL washing media. After a second round of centrifugation, the cells were resuspended in 5 mL of washing media and counted for further use.

### SHOCO seeding

Tumoroids and CAFs were harvested, and PBMCs were thawed following the protocols previously described. One 25 µL tumoroid dome was used for a total 100 µL SHOCO volume (1:4 dilution). Additionally, 3 × 10⁶ PBMCs (30000 PBMCs / µL) and 5 × 10⁵ CAFs (5000 CAFs / µL) were mixed with the tumoroids. This total volume generated 20x SHOCOs of 5 µL per well each, and could be scaled upwards depending on the experimental needs. The cell suspension containing all three populations was then centrifuged (3 min, 300 g), the supernatant was completely removed without disturbing the pellet, and finally, the cells were resuspended in 2 mg/mL Collagen I (5225-50ML, BIOMATRIX) and kept on ice until needed. Two protocols were employed over the course of this study to seed SHOCOs, depending on the throughput needs of the experiment: 1. Manual seeding of SHOCOs and 2. Semi-Automatic seeding of SHOCOs.

In preparation for seeding SHOCOs, 100 µL of dOil (HFE3500, 3M) + 0.5% Surfactant (dSURF, Fluigent) were added to each well in an ultra-low attachment U-bottom 96-well plate (MS-9096UZ, SBio). Five microliters of cell suspension in hydrogel were added to each well. Importantly, the gel was not allowed to touch the walls of the plate and had to be completely submerged in the oil. This allowed for a hydrogel + cell sphere to form and float right under the surface of the oil. The plate with the seeded SHOCOs was placed on a shaker (130 rpm) in the incubator for 20 minutes for gel polymerization. Once the collagen polymerized, the oil was carefully removed and pooled for further use. The plate was then left open under the hood to allow oil residues to evaporate. Afterward, 100 µL of SHOCO media (1:1 Tumoroid media + CAF media) was added to each well. The plate was then placed in an incubator on a shaker (130 rpm) for culturing.

For higher throughput seeding of SHOCOs, we used a dispenser robot (Integra Assist Plus). This method was employed when multiple 96-well plates were needed and is designed to allow full plate seeding and oil removal/washing of SHOCOs. The essential settings needed for this approach were the aspiration speed and adjustable height of the pipette tip. The robot prepared the plates with the necessary oil, after which it seeded the cell-hydrogel mix in 5 µL droplets per well under the surface of the oil - hence the need for adjustable height settings. The plates were then placed in the incubator for 20 minutes. Oil removal was performed by setting the robot to reach the bottom of the well and aspirate the oil at the lowest speed setting. The combination of low suction forces and the slippery water droplets ensured no SHOCOs were mistakenly removed. The robot then dispensed the culture media on top.

### SHOCO treatment

Fresh SHOCO cultures were maintained for 48 hours to allow for self-organization. Then, the media was replaced with media containing the required compounds, marking the start of treatment (day 0) - time points of interest were days 1, 3, and 5. To ensure continuous SHOCO viability, all samples, regardless of their intended treatment period, received a complete media refresh without any additional compounds on day 3. At these time points, the media was completely removed from the corresponding samples and frozen at -20°C for cytokine analysis. The SHOCOs were washed once with 100 µL per well of DPBS and transferred to a prepared 96-well plate with 100 µL per well of 4% PFA solution (J19943.K2, Thermo Scientific) for overnight fixation at 4°C. On the second day, the PFA was removed, and the samples were washed twice with 150 µL per sample of DPBS before adding 100 µL of DPBS to avoid drying and sealing the plate. Fixed samples were kept in the fridge at 4°C until needed.

### Hypoxia measurement^104^

Hypoxia levels in SHOCOs were measured using an EF5 Hypoxia Detection Kit (EF5-30A4M, Sigma-Aldrich). The EF5 stock solution (10 mM) was created following the manufacturer’s instructions and used at a working concentration of 50 µM. SHOCOs intended to be fixed at a certain time point were incubated for 4.5 hours with EF5 prior to fixation, allowing the compound to enter and tag hypoxic cells. The SHOCOs were then embedded in paraffin and marked using immunofluorescence antibodies as described below. As part of the antibody panel, anti-EF5 Alexa Fluor 488 (diluted 1:100) from the hypoxia kit was used to target the EF5 present in hypoxic cells. A hypoxic control was used with SHOCOs cultured in an incubator set at 5% O₂.

### PBMC infiltration assay

Tumoroids and CAFs were harvested, and PBMCs were thawed according to previously described protocols. Tumoroids (1:4 dilution), PBMCs (3000 PBMCs/µL), and CAFs (5000 CAFs/µL) were mixed, and the resulting cell suspension was centrifuged (3 min, 300 g). The supernatant was completely aspirated without disturbing the pellet, and the cells were resuspended in 2 mg/mL Collagen I (5225-50ML, BIOMATRIX) on ice. The hydrogel mixture was seeded at 5 µL to form SHOCOs or domes, followed by the addition of 100 µL of SHOCO media to each well and incubation on an orbital shaker (130 rpm) for 2 days. On day 2, the media was removed and replaced with fresh media containing PBMCs (1.3x10^5^ per well) and 0.1 µg/mL EpCAM TCB, then incubated under static conditions. After 3 days of treatment, the SHOCOs or domes were washed once with 100 µL per well of DPBS and transferred to a 96-well plate containing 100 µL per well of 4% PFA solution (J19943.K2, Thermo Scientific) for overnight fixation at 4°C. The following day, the PFA was removed, samples were washed twice with 150 µL per sample of DPBS, and 100 µL of DPBS was added to prevent drying before sealing the plate for subsequent Histogel embedding.

### SHOCO embedding in Histogel and Paraffin^105^

Solid Histogel (HG-4000-012, Epredia) was liquefied in the microwave for 25 seconds and poured onto the desired tissue microarray mold (48 samples) until full. The mold was then placed in the fridge at 4°C for 15 minutes. Once the Histogel solidified, it was removed from the mold and turned upside down, revealing cavities that allow for the organized embedding of multiple samples. Fixed SHOCOs were then placed individually in each cavity. Ten microliters of liquid Histogel were carefully added on top of each SHOCO to help "glue" it to the block and left to dry at room temperature for 1 minute before the block was filled with liquid Histogel, completely enclosing the SHOCOs inside. The samples were returned to the fridge for 15 minutes. Once the block containing the samples was solid, it was placed in a plastic cassette and then in a 10% Formalin (HT501128, Sigma Aldrich) solution in preparation for the dehydration and paraffin-embedding process that followed on the same day. The dehydration protocol was performed with a tissue processor (HistoCore Pearl, Leica). The samples were first treated with 10% Formalin (30 min), followed by sequential treatments with 70% ethanol (30 min and 1 hour), 80% ethanol (30 min), 96% ethanol (30 min and 1 hour), and 100% ethanol (1 hour and 45 min). The ethanol-treated samples were then treated with xylene (45 min, twice). Finally, the samples were treated with paraffin three times (45 min each) at 56°C. On the following day, the samples were embedded in liquid paraffin using metallic molds and fastened to the initial cassette, which acts as an anchoring point for the microtome. FFPE blocks were sectioned at a thickness of 4 µm and transferred onto IHC adhesive glass slides (TOMO, TOM-1190), which were then incubated overnight at 37°C.

### FFPE-slides multiplexed immunofluorescence (mIF) staining - Opal dyes

mIF staining was performed following an already published protocol^27,105^ using the Ventana Discovery Ultra automated tissue stainer (Roche Tissue Diagnostics, Tucson, AZ, USA). In a nutshell, FFPE slides were deparaffinized and subjected to antigen retrieval, followed by a blocking step. In a sequential process, a primary antibody, a corresponding secondary antibody, and then an Opal dye (480, 520, 570, 620, 690, or 780) were applied. Each cycle was followed by a denaturation and antibody neutralization step to prepare the samples for the next staining cycle. Lastly, samples were counterstained with 4’,6-Diamidino-2-phenylindole (DAPI, Roche), and slides were mounted manually using ProLong™ Gold Antifade Mountant (#P36930, Invitrogen). Full antibody specifications are provided in Table S2.

### Brightfield imaging

SHOCOs were imaged using a Nikon Ti-2 microscope, with one sample per U-bottom well and at 4x magnification. SHOCOs were then automatically detected using ImageJ software, and their sizes were compared to measure ongoing shrinking.

### Analysis of immunofluorescence images

Image analysis was conducted using self-trained Deep Net neural networks (DenseNet AI V2 Plugin) from HALO AI (Indica Labs). Separate neural networks were trained for each experiment to account for differences in staining intensity and background between immunofluorescence staining and scanning runs. Training regions were manually annotated across multiple SHOCOs and tissue sections, and segmentation outputs were visually inspected and iteratively refined. Quality control also included sections and SHOCOs that were not used for manual annotation, ensuring that classifier performance was maintained across biological replicates and different sectioning depths. For SHOCOs treated with EpCAM-TCB (**Figure 3**), each SHOCO was segmented individually, and the average IF signal per SHOCO was used for analysis. In the case of SHOCOs with different CAF concentrations and CEACAM5-TCB treatment (**Figure 4**), epithelial niches and stromal regions (around 20-30 objects in total) were manually selected and used as training data for the neural network (**Figure S3**). Furthermore, the same software was used to detect single cells by DAPI expression and phenotype them as CD4 or CD8 T cells, depending on the signal measured in those cells. Combining these processes and performing TMA batch analysis, we measured signals per cell (e.g., cleaved caspase-3/7) specifically in different regions, as well as analyzed the images by single-cell phenotyping of T cells for each region inside individual SHOCOs.

Analysis of hypoxia levels by different layers (**Figure 1H**) was performed following a different workflow. First, SHOCOs were segmented individually, resulting in their outlines forming annotations. Using these annotation lines delimiting the SHOCO edge, further annotations were automatically drawn in 100 µm steps towards the center of the object, forming three different layers (center, middle, outer). Hypoxia levels were then measured per cell and averaged per layer.

### SHOCO digestion for flow cytometry

To prepare a single-cell suspension for flow cytometry, SHOCOs were resuspended in 5mL of digestion cocktail consisting of RPMI-1640 medium supplemented with glutamax (Gibco, 61870-010), 5% heat inactivated FBS (Gibco, A5209401), 10 mmol/L Hepes (Gibco, 15630-056), 40U/mL DNase-I (Roche, 10104159001), 2mg/mL Collagenase-D (Roche, 11088858001) and 1mg/mL trypsin inhibitor (Sigma, T9128). Samples were added into pre-coated (0.1 % BSA in PBS) gentle MACS C Tubes (Milteny, 130-093-237) and were mechanistically dissociated using the standard 37C_h_TDK_3 program on a gentleMACS dissociator with heaters, according to the manufacturer’s instructions. Following dissociation, the cell suspension was diluted with complete RPMI-1640 medium, passed through a 70µm cell strainer (Miltenyi Biotec, 130-095-823) to remove debris, and centrifuged to pellet the cells for downstream flow cytometry staining.

### Flow cytometry staining

For extracellular staining ten domes or SHOCOs were pooled and digested to single cells, Fc receptors were blocked by incubating cells with a 2X concentration of Human TruStain FcX (BioLegend, 422302) in FACS buffer for 15 min at 4°C. Without washing, cells were stained for 30 min at 4°C with a 2X cocktail of surface antibodies diluted in a mixture of BD Horizon Brilliant Stain Buffer (BD Biosciences, Cat# 566349) and FACS buffer (1:5 ratio). Cells were then washed with PBS and incubated with a fixable viability dye (Live/Dead) for 15 min at 4°C to exclude dead cells. For intracellular staining cells were washed with permeabilization buffer and fixed using the Foxp3/Transcription Factor Staining Buffer Set (Thermo Fisher Scientific, Cat# 00-5521-00) for 30 min at room temperature, following the manufacturer’s protocol. After fixation, cells were washed in permeabilization buffer and incubated with intracellular antibodies for 30 min at 4°C. Finally, cells were rinsed with permeabilization buffer and passed through a cell-strainer cap prior to sample acquisition on a Cytek Aurora full-spectrum flow cytometer (Cytek Biosciences). All resulting FCS files were analyzed using FlowJo software (v11, BD Biosciences). Full antibody specifications are provided in Table S2.

### Cytokine Analysis

Cytokine analysis was performed on frozen supernatants using a bead-based multiplex kit, following the manufacturer’s protocol. For EpCAM-TCB treatments (**Figure 3**), a standard Human CD8/NK-cell panel (741187, BioLegend) was used, while a custom panel consisting of TNFα, TGF-β1, IL-1β, CXCL10, IL-2, CCL2, IL-10, sFas, CXCL8, sFasL, IL-6, IFNγ, and Granzyme B (GZMB) was employed for the CEACAM5-TCB treated samples (**Figure 4**).

Briefly, 1:10 diluted supernatants were mixed (in equal amounts) with cytokine-binding beads and assay buffer (provided in the kit), then placed on a shaker (400 rpm) for 2 hours at room temperature. The beads were subsequently washed and incubated with detecting antibodies (provided in the kit) for 1 hour, followed by an additional 30-minute incubation with SA-PE antibodies (provided in the kit). After this, the beads were washed to remove excess antibodies, suspended in the washing buffer (provided in the kit), and acquired using a flow cytometer (Cytek Aurora) using the manufacturer’s recommended experiment setup. The raw data were processed to obtain absolute cytokine concentrations using the provided software (www.legendplex.qognit.com).

### Imaging Mass Cytometry (IMC) data acquisition^106^

#### Antibody labelling

Antibody conjugations to lanthanide metals via a polymer linker were performed using the MaxPAR antibody labeling kit (Standard BioTools) according to manufacturer instructions. The yield of the antibody after this process was quantified using a Nanodrop One Spectrophotometer (Thermo Scientific), then diluted with Tris Antibody Stabilizing buffer (Candor Bioscience) for storage at 4°C. All conjugated antibodies were titrated on a set of reference cell lines to determine the optimal staining concentrations for each target prior to panel preparation and sample staining.

#### IMC sample staining

FFPE sample dewaxing, antigen retrieval and antibody staining was performed on a Leica Bond-RXm autostainer (Biosystems). A first baking step at 60°C for 30 min was followed by dewaxing at 72°C for 30min and tissue rehydration in an ethanol gradient. Epitope retrieval was then performed at 95°C for 40 min at a pH of 9 using BOND epitope retrieval solution 2 (Biosystems). After washing steps, the sample was blocked with Novocastra protein block (Biosystems) for 30 min at room temperature, followed by staining with the antibody panel for 3.5 h, also at room temperature. DNA staining was then performed using an iridium intercalator at 0.5µM for 5 minutes, followed by washing. After a quick dip in deionized water, the sample slides were dried with pressurized air and stored at room temperature until acquisition.

#### IMC data acquisition

IMC images were acquired using a CyTOF XT Mass Cytometer coupled to a Hyperion XTi imaging system (Standard Biotools). Once the instrument was calibrated, samples were ablated with a UV laser frequency of 800Hz. A panel-specific spillover slide was acquired and used to compensate for potential spillover between channels as previously described (IMC1).

### IMC data processing

#### Segmentation

We use a pretrained neural network for segmentation - DeepCell^107^. The input to the network consists of a nuclear and a cytoplasm channel. The used IMC panel does not contain a general cytoplasm marker. To construct a cytoplasm channel that contains signals for all major cell types, we pool information from multiple channels. We first normalize each channel such that all pixel values are between 0 and 1 to account for differences in signal intensity between the different markers. Next, we calculate the general cytoplasm channel by taking the maximum pixel value over the following channels for each pixel: 115In_SMA, 141Pr_CA9, 143Nd_panK, 146Nd_CD45, 147Sm_CD31, 160Gd_GPA33, 176Yb_Vimentin, 209Bi_K8-18. The output of the DeepCell classification network consists of nuclear and whole cell predictions. Since these predictions are not necessarily consistent (e.g. one whole cell may have multiple nuclei), we perform a postprocessing step in which we make sure that every whole cell contains exactly one nucleus. We achieve this by 1) deleting all whole cells that contain either more than one or zero nuclei and 2) generating whole cells for nuclei that do not lie within a whole cell by dilating the nucleus by 2 mm.

#### Single cell analysis

For each cell and channel, we quantify marker expression by averaging the signal over all of the cell’s pixels. Additionally, we perform an interaction frequency analysis by identifying all pairs of cells that are less than 5 µm apart. We calculate the mean interaction count for target cells by averaging the interactions across all replicates in each system to determine the interaction frequency. Additionally, the interaction frequency analysis was performed on further 52 samples acquired from AMSBio and processed as described. The antibody panel used on these samples as part of a different study had an overlap of 29/40 including all markers applied in phenotyping and clustering.

#### Cell type classification

To classify the cell types of individual cells we trained an image based neural network by customizing the approach of CellSighter^108^ to our data. To train this network, we first generated training data by identifying cells with unambiguous marker expression through manual gating using the expression values of the markers displayed. We next trained and tested the network using an 80/20 split. Finally, we used the trained network to predict the cell types of all cells.

## Statistical Analysis

Statistical analysis details are provided in the description of each figure. The number of donors is indicated as ‘N=x’, whereas the number of independent scaffolds or technical replicates is referred to as ‘n=x’ in the respective legend text. P values indicating statistical significance are marked as follow: P ≥ 0.05 (ns); P < 0.05 (*); P < 0.01 (**); P < 0.001 (***); P < 0.0001 (****). All graphs and statistical analyses were generated using GraphPad Prism 10, R or Jupyter Notebook (Python).

For the neighborhood analysis of the different myeloid populations (**Figures 3J** and **3L**), statistical significance was assessed through permutation distribution (**Figure S3H**).

### Single-cell dissociation of the scaffolds for scRNA-seq

Prior to dissociation, surfaces were cleaned and all tubes were coated with a wash buffer containing HBSS -Ca+/ -Mg2+ (14175095, Gibco) + 1% BSA (130-091-376, Miltenyi Biotec) on ice for at least 15 min.

To dissociate SHOCO cultures, structures were transferred to pre-coated tubes containing a digestion cocktail composed of RPMI-1640 medium with GlutaMAX (Gibco, 61870-010), 5% heat-inactivated FBS (Gibco, A5209401), 10 mM HEPES (Gibco, 15630-056), 40 U/mL DNase I (Roche, 10104159001), 2 mg/mL Collagenase D (Roche, 11088858001), and 1 mg/mL trypsin inhibitor (Sigma-Aldrich, T9128). Enzymatic digestion was performed on a thermoshaker (Eppendorf ThermoMixer C) at 37°C and 1,000 rpm using alternating 3–5 min incubation cycles interspersed with vigorous mechanical trituration until complete tissue disaggregation was achieved. Samples were pelleted by centrifugation (500 g, 4°C, 5 min), washed with PBS, and further incubated TrypLE Express (Thermo Fisher Scientific, 12604013) under identical thermoshaker conditions (37°C, 1,000 rpm) with periodic resuspension to yield single cells. The reaction was quenched with culture medium, washed with PBS, and pooled by patient line.

For rBM-dome co-cultures, matrix structures were washed once with PBS and incubated with Cell Recovery Solution (Corning, 354253) for 30 min at 4°C to release embedded cells. Isolated cells were pooled, pelleted (500 g, 4°C), subjected to TrypLE Express dissociation as described above, and washed with PBS.

For 2D monocultures, CAFs were washed with PBS, incubated with TrypLE Express for 5 min at 37°C, and pelleted by centrifugation (500 g, 4°C).

Single-cell suspensions were incubated with Zombie UV Fixable Viability Dye (1:1000, 423107, BioLegend) and FcR Blocking Reagent (1:100, 130-059-901, BioLegend) in PBS for 30 min at 4°C protected from light. Following centrifugation (500 g, 4°C) and a wash step with FACS buffer (PBS supplemented with FBS), cells were incubated with anti-EpCAM-Alexa Fluor 700 antibody (1:100, 56-5791-82, Invitrogen) in FACS buffer on ice for 20 min. After a final wash, cells were resuspended in a wash buffer supplemented with a ROCK inhibitor (Y-27632). Single cells from SHOCO and rBM-dome cultures were sorted into EpCAM– and EpCAM+ populations using a Cytek Aurora CS cell sorter, whereas 2D CAF monocultures were processed directly for downstream analysis.

### scRNA-seq library preparation and read alignment of single cell genomic data

Following sorting, cells were counted on a Cellaca MX High-Throughput Automated Cell Counter (Nexcelom Bioscience) using an acridine orange/propidium iodide stain and mixed at a 30:70 ratio of epithelial cells to CAFs for each condition.

Single-cell gene expression libraries were prepared using the GEM-X Universal 3’ v4 Gene Expression assay (10x Genomics) according to Chromium GEM-X Single Cell 3’ v4 Gene Expression User Guide (CG000731, Rev B). A total of 20,000 cells were targeted per sample. Libraries were sequenced on the NovaSeq X Plus (Illumina) with a read configuration of 28-10-10- 90, targeting 35,000 reads per cell. Raw sequencing data underwent base calling using bcl-convert (v4.3.13, Illumina). The quality of the resulting FASTQ files was assessed using FastQC (v0.12.1). FASTQ files were processed with Cell Ranger (v9.0.1, 10x Genomics), and reads were aligned to the human reference GRCh38-2024-A (10x Genomics).

### Single-cell RNA-seq analysis

Filtered feature-barcode matrices were imported into Seurat, and individual sample objects were merged for downstream analysis. High-quality cells with more than 1,000 detected genes and less than 10% mitochondrial transcripts were retained. Data were normalized using NormalizeData, 2,000 highly variable features were identified using the vst method, and data were scaled. Principal component analysis (PCA) was performed using variable features, followed by nearest-neighbor graph construction, clustering, and UMAP visualization using the first 20 principal components. CAFs and epithelial cells were identified based on the expression of COL1A1 and EPCAM, respectively, separated into two objects and reprocessed using the same workflow described above. Batch correction across samples was performed using Harmony integration^109^. The resulting graph was used for ‘FindClusters’ with the Louvain algorithm at a resolution of 0.3 for CAFs and 0.5 for epithelial cells.

To identify cell-type-specific transcriptional changes, differential gene expression (DEG) analysis was conducted using the FindMarkers function in the Seurat R package. A gene was considered differentially expressed if it was detected in a minimum of 10% of cells in either condition (min.pct = 0.1) and met the significance thresholds of an adjusted p-value < 0.05 and an absolute log2 fold change > 1. Subsequently, a functional Over-Representation Analysis (ORA) was conducted specifically on the list of genes significantly upregulated in co-culture (log2 fold change > 1). This analysis was performed in parallel against three major databases using the clusterProfiler^110^(4.12.6) R package: the Gene Ontology (GO) biological processes, KEGG pathways, and the MSigDB Hallmark gene sets. For all tests, a custom background consisting of all detected genes in the dataset was used as the statistical universe. Significantly enriched terms were defined by a p-value < 0.05 and a q-value < 0.2. GO results were additionally simplified to reduce semantic redundancy. Published in vivo CRC fibroblast gene programs from Pelka et al.^31^ were used to score CAFs from the three models using the AddModuleScore_UCell function.

For consensus non-negative matrix factorization (cNMF) analysis^56^, the CAF object was downsampled to 10,000 cells per sample (n=60,000 total cells). cNMF was run with a number of components (k) from 5 to 20, with 200 iterations and the 2,000 most variable genes detected in Seurat. We selected k=14 as the preferred solution considering stability and error and removed outliers with Euclidean distance > 0.1. cNMF program scores within cells were normalized to sum to 1. Pathway enrichment analysis was performed on the top 50 genes for each cNMF program using clusterProfiler with the Gene Ontology Biological Process and MSigDB Hallmark gene sets. To elucidate the upstream regulators governing intercellular crosstalk, a computational analysis was performed using NicheNet^57^ (2.2.1). The model was designed to identify ligands expressed by CAFs populations capable of inducing the transcriptional response observed in receiver populations (epithelial and CAFs) comparing rBM-dome to SHOCOs. This transcriptional response was defined as the set of differentially expressed genes (adjusted p-value < 0.05; |log2 fold change| > 0.25) between these conditions. The regulatory potential of each ligand was inferred by quantifying the enrichment of its a priori-defined target genes within this differential expression signature, with ligand activity scored by the Area Under the Precision-Recall curve (AUPR). The top 30 ligands with the highest AUPR scores were prioritized as the most probable upstream regulators for subsequent characterization of their expression and predicted target gene networks.

---

**Figure S1.**
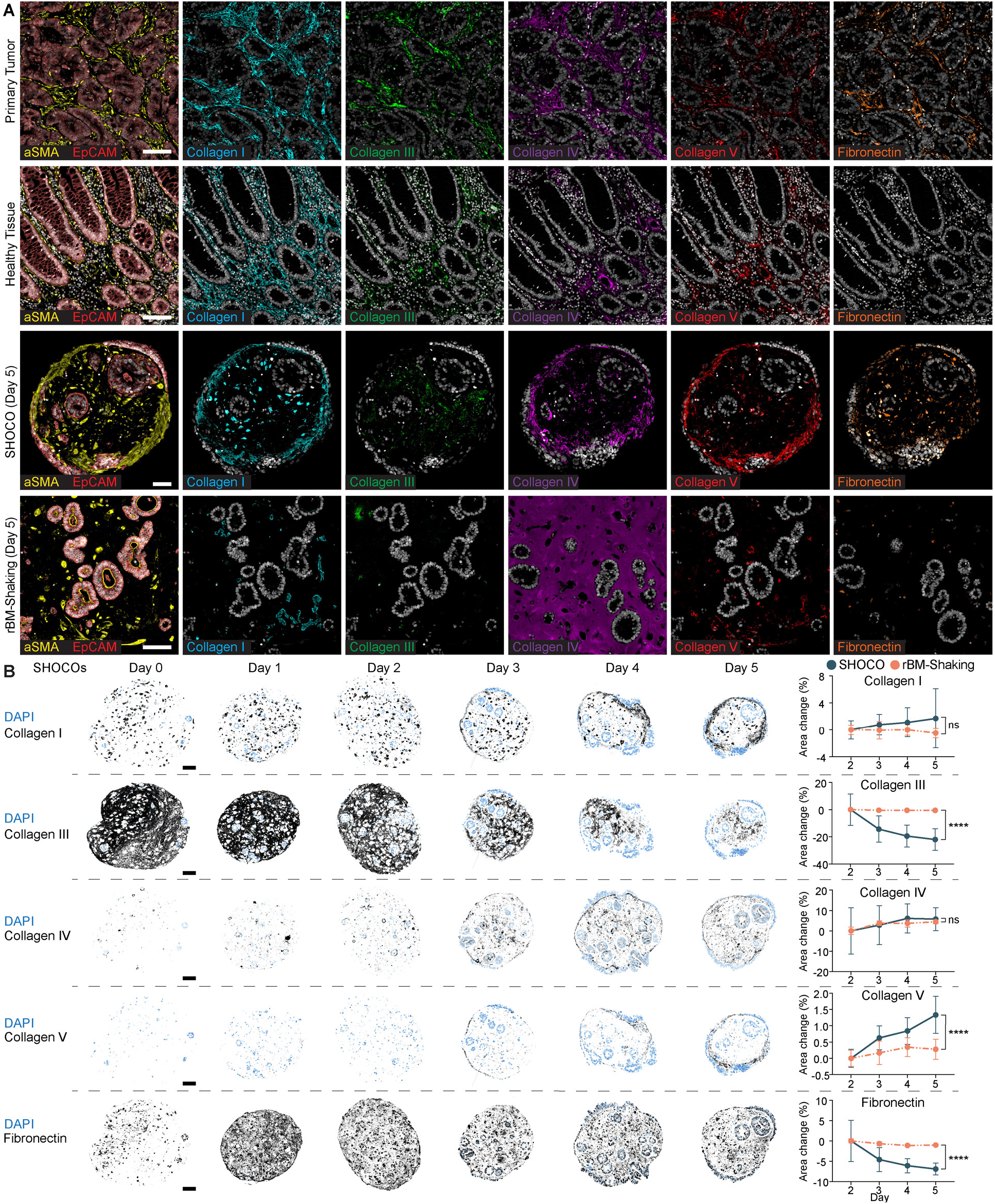
SHOCOs establish a complex tumor-like ECM that undergoes dynamic remodeling over time compared with rBM-based cultures. **(A)** Multiplex immunofluorescence images of Col1A2, Col3A1, Col4A1, collagen V fibrils, and fibronectin fibrils in primary tumor, healthy tissue, day 5 SHOCOs, and day 5 rBM-Shaking cultures. *a*-SMA and EpCAM mark stromal and epithelial compartments, respectively. **(B)** Spatial distribution of collagen I, collagen III, collagen IV, collagen V, and fibronectin in SHOCOs from days 0–5. Quantification shows the change in ECM-positive area from days 2–5 in SHOCOs compared with rBM-Shaking cultures. *N* = 2 SHOCO donors (Donor 1, Donor 5); *n* = 4 SHOCOs/donor; 2 FFPE sections/SHOCO. *N* = 1 rBM-Shaking donors (Donor 1); *n* = 4 cultures/donor; 2 FFPE sections/culture. Data are presented as mean ± SD and were analysed by ordinary two-way ANOVA (culture system × time), followed by Šídák’s multiple- comparisons test comparing culture systems at each time point (single pooled variance). Only the comparison at day 5 is annotated. P ≥ 0.05 (ns); P < 0.05 (*); P < 0.01 (**); P < 0.001 (***); P < 0.0001 (****). Scale bars are 100 μm (white and black).

**Figure S2.**
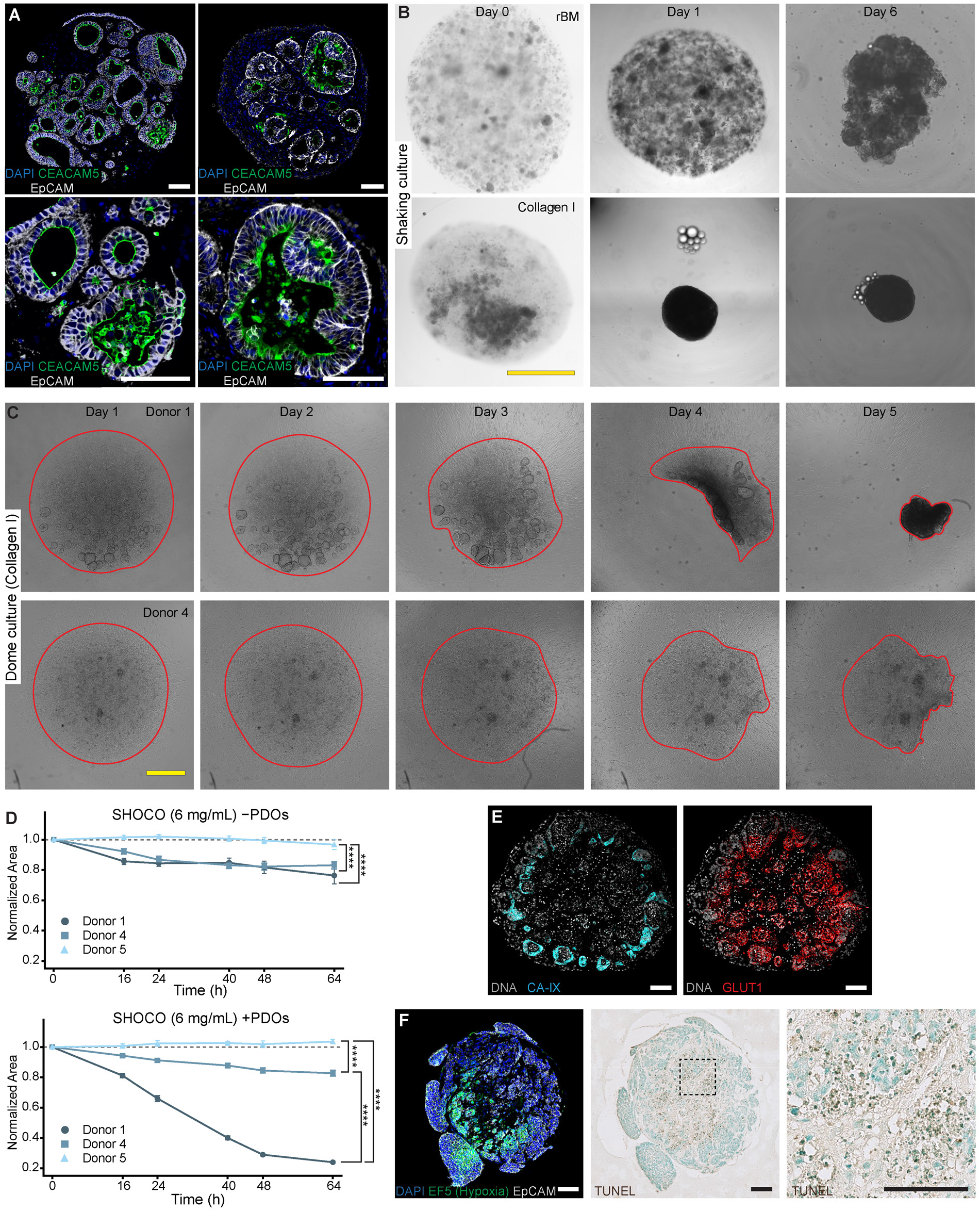
Donor-specific CAF-mediated collagen compaction drives SHOCO shrinkage and creates a dense microenvironment with hypoxic and necrotic cores. (A) Multiplex immunofluorescence images of day 5 SHOCOs from Donor 1 and Donor 5 showing apical CEACAM5 localization within EpCAM-positive epithelial structures. (B) Brightfield images of SHOCOs (Donor 1) and rBM-Shaking cultures (Donor 1) at days 0, 1, and 6. (C) Brightfield images of Col-Dome cultures from Donor 1 and Donor 4 over 5 days. Dome boundaries are outlined in red. (D) Quantification of matrix compaction in SHOCOs generated in 6 mg/mL collagen over 64 h. Cultures contained either CAFs alone or CAFs together with matched PDOs. Area was normalized to the corresponding measurement at 0 h. *N* = 3 (Donor 1, Donor 4, Donor 5); *n* = 11/14/12 SHOCOs without PDOs and 15/14/14 with PDOs. Data is shown as mean and analysed by Complete-time- course AUCs were compared using two-sided permutation tests with Benjamini–Hochberg FDR correction. **(E)** Spatial mass spectrometry images of a day 5 SHOCO showing DNA (gray) with carbonic anhydrase IX (CAIX; cyan) or GLUT1 (red). *N* = 1 (Donor 4); **(F)** Serial sections of a day 4 SHOCO (Donor 7) showing EF5- positive hypoxic regions and TUNEL staining. Right, higher-magnification image of the indicated TUNEL- positive region. P ≥ 0.05 (ns); P < 0.05 (*); P < 0.01 (**); P < 0.001 (***); P < 0.0001 (****). Scale bars are 100 μm (white and black) and 1 mm (yellow).

**Figure S3.**
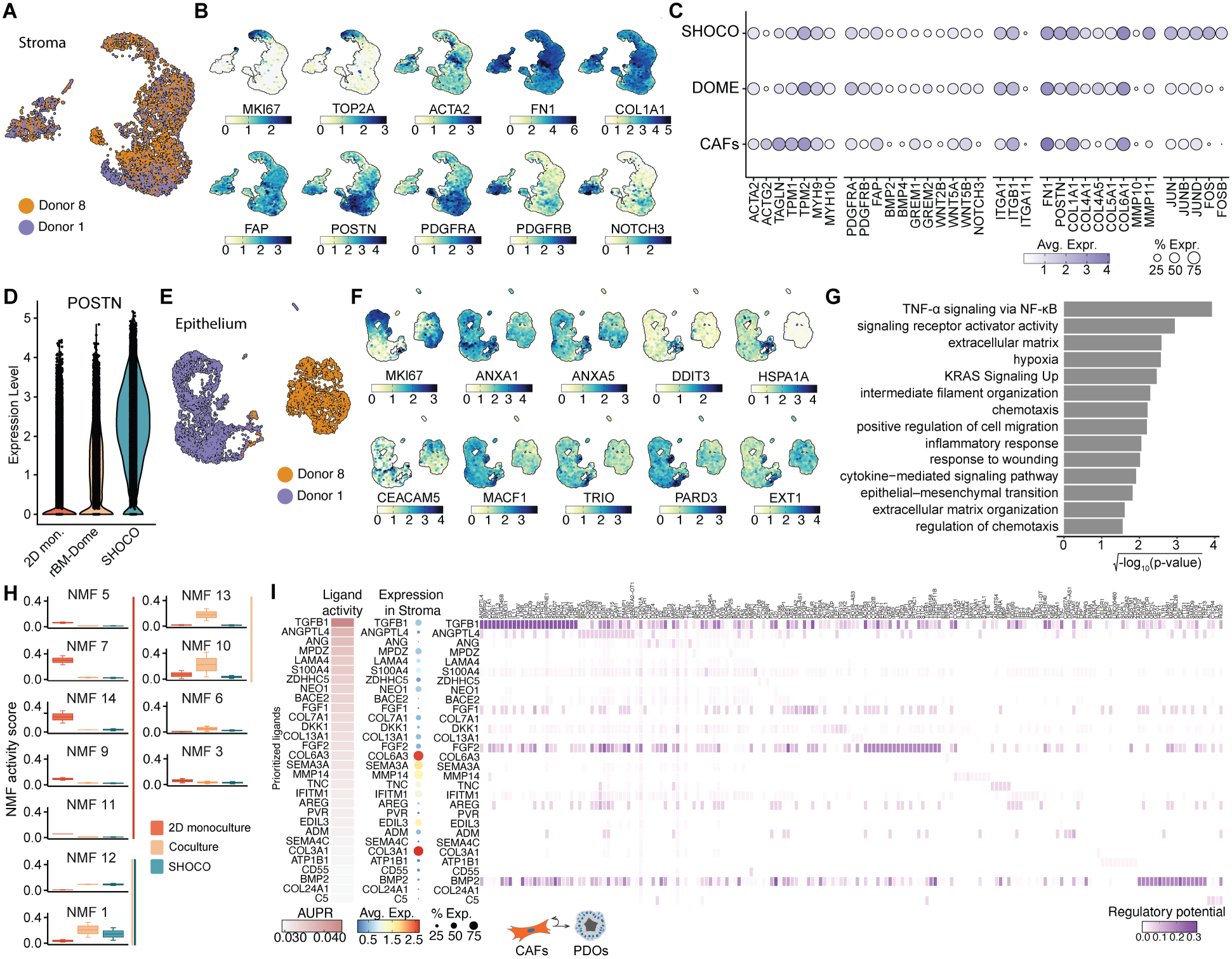
**SHOCOs induce distinct stromal and epithelial transcriptional programs revealed by single- cell and ligand–target analyses**. **(A)** Integrated UMAP embedding of scRNA-seq transcriptome data from stromal cells isolated from CAF monoculture, rBM-Dome culture and SHOCO, highlighting different donors. *N* = 2 (Donor 1, Donor 8). **(B)** UMAP expression plots of canonical genes highly expressed in the cell clusters. **(C)** Dot plot highlighting canonical marker gene expression across different culture formats. **(D)** Violin plot comparing the expression of POSTN in stomal cell from CAF monoculture, rBM-Dome culture and SHOCO. **(E)** Integrated UMAP embedding of scRNA-seq transcriptome data from epithelial cells isolated from rBM- Dome cultures or from SHOCOs, highlighting different donors. *N* = 2 (Donor 1, Donor 8). **(F)** UMAP expression plots of canonical CRC genes highly expressed in the cell clusters. **(G)** Enrichment analysis (GO, KEGG, and MSigDB Hallmarks) for genes upregulated in epithelial cells under different culture conditions (Table S1). Displayed pathways were manually selected for relevance. **(H)** Comparison of activity scores for cNMF- derived stromal gene-expression programs across culture systems. Data are shown as box-and-whisker plots with median as center line. **(I)** Heatmap of top-ranked stroma ligands prioritized by their predicted regulatory potential in SHOCOs. Ligands were ranked based on their potential to regulate DEGs upregulated in SHOCOs in the target cell types (stroma and epithelium). Dot plot of ligands expression in the stroma compartment. NicheNet’s ligand–target matrix denoting the regulatory potential between colonocyte-ligands (top) and fibroblast-ligands (bottom) and target genes.

**Figure S4.**
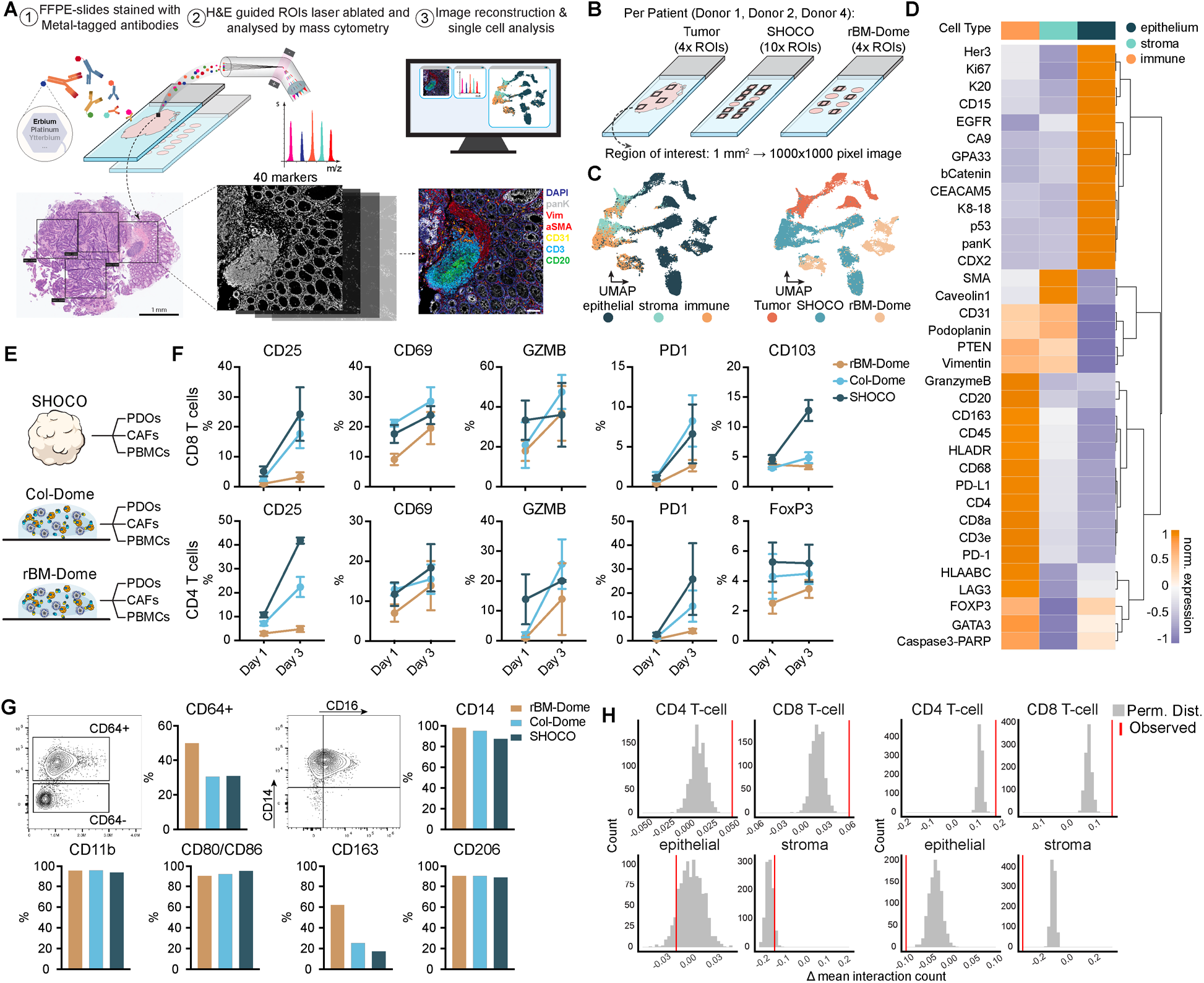
SHOCOs induce higher baseline T-cell activation than rBM-based cultures and provide an environment that supports monocyte-to-macrophage differentiation. (A) Schematic of the imaging mass cytometry (IMC) workflow. (B) Sampling strategy for primary tumors, SHOCOs, and rBM-Dome cultures from tumor Donor 1, Donor 2 and Donor 4, and allogeneic PBMC donor IC02. Four ROIs were analyzed per primary tumor and dome culture and ten ROIs per SHOCO. Each ROI covered 1 mm² and generated a 1,000 × 1,000- pixel image. (C) UMAPs of IMC-profiled cells colored by epithelial, stromal, and immune populations or by culture system. (D) Heatmap of normalized expression of the 40 IMC markers across epithelial, stromal, and immune populations. (E) Systems compared by flow cytometry: SHOCOs, Col-Dome or rBM-Dome cultures, each containing PDOs, CAFs, and PBMCs. *N* = 1 tumor donor (Donor 1) and 3 allogeneic PBMC donors (IC04, IC08, IC09); *n* = 10 replicates/condition. (F) Quantification of CD25, CD69, granzyme B, and PD-1 expression in CD8 and CD4 T cells at days 3 and 5, together with CD103 in CD8 T cells and FOXP3 in CD4 T cells. Data are presented as mean ± SD. (G) Flow cytometry phenotyping of myeloid cells at day 2 showing differentiation toward a macrophage-like phenotype across all cultures. Flow cytometry plots show CD64 expression and CD14/CD16 distribution; barplots quantify CD64, CD14, CD11b, CD80/CD86, CD163, and CD206. *N* = 1 tumor donor (Donor 1) and 1 allogeneic PBMC donor (IC10). (H) Permutation distributions for the interaction-frequency analyses shown in Figures 3J and 3L. Gray distributions indicate permuted interaction counts and red lines indicate the observed values.

**Figure S5.**
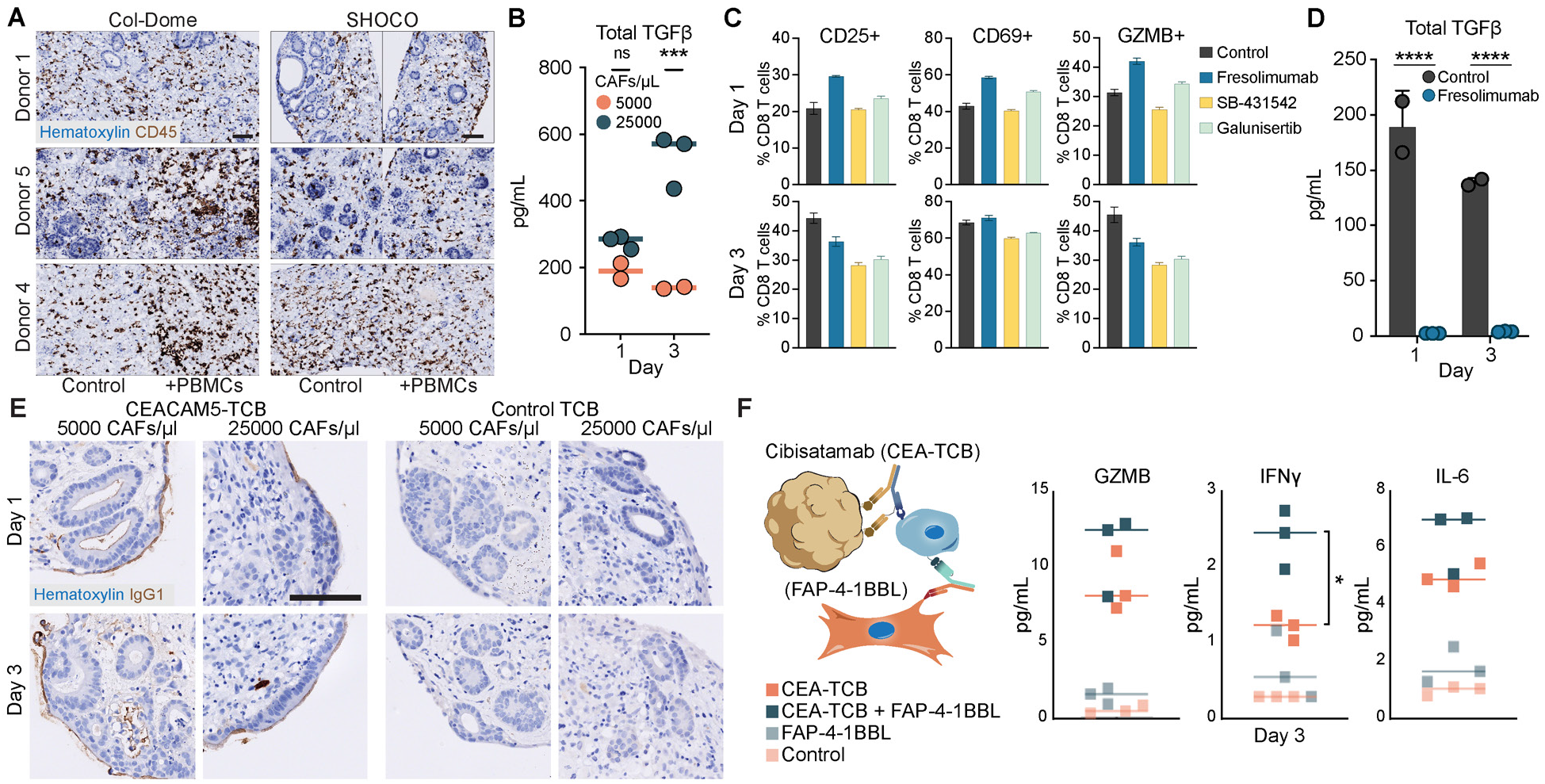
SHOCOs enable mechanistic interrogation of stromal barriers and immunoregulatory signals shaping TCB responses. **(A)** Representative CD45 IHC images of Col-Dome cultures and SHOCOs cultured with PBMCs in the supernatant or without (Control) and treated with EpCAM-TCB. Images were acquired on day 3. *N* = 3 tumor donors (Donor 1, Donor 4, Donor 5) and 1 allogeneic PBMC donor (IC07). **(B)** Total TGF-β levels in SHOCOs generated at low (5000 CAFs/µL) or high (25000 CAFs/µL) stromal density on days 1 and 3. Data are shown as individual replicates with median and were analysed by ordinary two-way ANOVA (stromal density × time), followed by Šídák’s multiple-comparisons test (single pooled variance). **(C)** CD25, CD69, and GZMB expression in CD8 T cells on days 1 and 3 following CEACAM5-TCB activation. SHOCOs were additionally treated with fresolimumab, SB-431542, galunisertib, or left untreated (Control). Data are presented as mean ± SD. **(D)** Total TGF-β levels in untreated and fresolimumab-treated SHOCOs on days 1 and 3. Data are shown as mean ± SD and were analysed by ordinary two-way ANOVA (treatment × time), followed by Šídák’s multiple-comparisons test (single pooled variance). **(E)** Representative anti-IgG1 IHC images of SHOCOs generated at low (5000 CAFs/µL) or high (25000 CAFs/µL) stromal density on days 1 and 3, with or without CEACAM5-TCB treatment. Samples used in (B-E) are *N* = 1 tumor donor (Donor 1) and 1 allogeneic PBMC donor (IC07); *n* = 10-12 SHOCOs per condition. **(F)** Illustration of the mechanism of action of cibisatamab (CEA-TCB) in combination with FAP-4-1BBL, together with day-3 measurements of GZMB, IFN-γ, and IL-6 following treatment with CEA-TCB or a control TCB, each with or without FAP-4-1BBL. The control TCB lacks a cognate antigen in the culture, leaving its masked anti-CD3 arm unavailable for T- cell receptor binding. *N* = 1 tumor donor (Donor 12); *n* = 3 SHOCOs/condition. Data were analysed by ordinary one-way ANOVA followed by Tukey’s multiple-comparisons test (single pooled variance). P ≥ 0.05 (ns); P < 0.05 (*); P < 0.01 (**); P < 0.001 (***); P < 0.0001 (****). Scale bars, 100 μm.

**Table S1.** Single cell sequencing data. (**A)** Top 40 cell type marker genes of stromal cells from Figure 2D; **(B)** Enrichment analysis (GO, KEGG, and MSigDB Hallmarks) for genes upregulated in stromal cells under different culture conditions for Figure 2E; **(C)** Gene programs from Pelka et al. used in Figure 2H and Figures S3H; **(D)** Pathway enrichment (GO Biological Process and MSigDB Hallmark) for the SHOCO-enriched cNMF program analyzed using clusterProfiler shown Figure 2;**. (E)** Enrichment analysis (GO, KEGG, and MSigDB Hallmarks) for genes upregulated in epithelial cells under different culture conditions for Figure 2E;

**Table S2.** Key resource table: **(A)** Primary human CRC patient donor metadata. **(B)** Antibody panel details.

## References

1. Herbst, R.S., Giaccone, G., Marinis, F.D., Reinmuth, N., Vergnenegre, A., Barrios, C.H., Morise, M., Felip, E., Andric, Z., Geater, S., et al. (2020). Atezolizumab for First-Line Treatment of PD-L1–Selected Patients with NSCLC. New England Journal of Medicine 383, 1328–1339. 10.1056/nejmoa1917346.

2. Reck, M., Rodríguez-Abreu, D., Robinson, A.G., Hui, R., Csőszi, T., Fülöp, A., Gottfried, M., Peled, N., Tafreshi, A., Cuffe, S., et al. (2016). Pembrolizumab versus Chemotherapy for PD-L1–Positive Non–Small- Cell Lung Cancer. New England Journal of Medicine 375, 1823–1833. 10.1056/nejmoa1606774.

3. Neelapu, S.S., Locke, F.L., Bartlett, N.L., Lekakis, L.J., Miklos, D.B., Jacobson, C.A., Braunschweig, I., Oluwole, O.O., Siddiqi, T., Lin, Y., et al. (2017). Axicabtagene Ciloleucel CAR T-Cell Therapy in Refractory Large B-Cell Lymphoma. New England Journal of Medicine 377, 2531–2544. 10.1056/nejmoa1707447.

4. Abramson, J.S., Palomba, M.L., Gordon, L.I., Lunning, M.A., Wang, M., Arnason, J., Mehta, A., Purev, E., Maloney, D.G., Andreadis, C., et al. (2020). Lisocabtagene maraleucel for patients with relapsed or refractory large B-cell lymphomas (TRANSCEND NHL 001): a multicentre seamless design study. Lancet 396, 839–852. 10.1016/s0140-6736(20)31366-0.

5. Wong, C.H., Siah, K.W., and Lo, A.W. (2019). Estimation of clinical trial success rates and related parameters. Biostatistics 20, 273–286. 10.1093/biostatistics/kxx069.

6. Haslam, A., Olivier, T., Powell, K., Tuia, J., and Prasad, V. (2023). Eventual success rate and predictors of success for oncology drugs tested in phase I trials. Int. J. Cancer 152, 276–282. 10.1002/ijc.34181.

7. Hegde, P.S., and Chen, D.S. (2020). Top 10 Challenges in Cancer Immunotherapy. Immunity 52, 17–35. 10.1016/j.immuni.2019.12.011.

8. Loewa, A., Feng, J.J., and Hedtrich, S. (2023). Human disease models in drug development. Nature Reviews Bioengineering 1, 545–559. 10.1038/s44222-023-00063-3.

9. Kim, J., Koo, B.-K., and Knoblich, J.A. (2020). Human organoids: model systems for human biology and medicine. Nature Reviews Molecular Cell Biology 21, 571–584. 10.1038/s41580-020-0259-3.

10. Day, C.-P., Merlino, G., and Terry (2015). Preclinical Mouse Cancer Models: A Maze of Opportunities and Challenges. Cell 163, 39–53. 10.1016/j.cell.2015.08.068.

11. Mak, I.W., Evaniew, N., and Ghert, M. (2014). Lost in translation: animal models and clinical trials in cancer treatment. Am J Transl Res 6, 114–118.

12. Bareham, B., Georgakopoulos, N., Matas-Céspedes, A., Curran, M., and Saeb-Parsy, K. (2021). Modeling human tumor-immune environments in vivo for the preclinical assessment of immunotherapies. Cancer Immunology, Immunotherapy 70, 2737–2750. 10.1007/s00262-021-02897-5.

13. Tian, H., Lyu, Y., Yang, Y.-G., and Hu, Z. (2020). Humanized Rodent Models for Cancer Research. Frontiers in Oncology 10. 10.3389/fonc.2020.01696.

14. Sato, T., and Clevers, H. (2013). Growing self-organizing mini-guts from a single intestinal stem cell: mechanism and applications. Science 340, 1190–1194. 10.1126/science.1234852.

15. Ooft, S.N., Weeber, F., Dijkstra, K.K., McLean, C.M., Kaing, S., van Werkhoven, E., Schipper, L., Hoes, L., Vis, D.J., van de Haar, J., et al. (2019). Patient-derived organoids can predict response to chemotherapy in metastatic colorectal cancer patients. Sci Transl Med 11. 10.1126/scitranslmed.aay2574.

16. Vlachogiannis, G., Hedayat, S., Vatsiou, A., Jamin, Y., Fernández-Mateos, J., Khan, K., Lampis, A., Eason, K., Huntingford, I., Burke, R., et al. (2018). Patient-derived organoids model treatment response of metastatic gastrointestinal cancers. Science 359, 920–926. 10.1126/science.aao2774.

17. van de Wetering, M., Francies, H.E., Francis, J.M., Bounova, G., Iorio, F., Pronk, A., van Houdt, W., van Gorp, J., Taylor-Weiner, A., Kester, L., et al. (2015). Prospective Derivation of a Living Organoid Biobank of Colorectal Cancer Patients. Cell 161, 933–945. 10.1016/j.cell.2015.03.053.

18. Ganesh, K., Wu, C., O’Rourke, K.P., Szeglin, B.C., Zheng, Y., Sauvé, C.-E.G., Adileh, M., Wasserman, I., Marco, M.R., Kim, A.S., et al. (2019). A rectal cancer organoid platform to study individual responses to chemoradiation. Nat. Med. 25, 1607–1614. 10.1038/s41591-019-0584-2.

19. Kim, E., Choi, S., Kang, B., Kong, J., Kim, Y., Yoon, W.H., Lee, H.R., Kim, S., Kim, H.M., Lee, H., et al. (2020). Creation of bladder assembloids mimicking tissue regeneration and cancer. Nature 588, 664–669. 10.1038/s41586-020-3034-x.

20. Sharpe, B.P., Nazlamova, L.A., Tse, C., Johnston, D.A., Thomas, J., Blyth, R., Pickering, O.J., Grace, B., Harrington, J., Rajak, R., et al. (2024). Patient-derived tumor organoid and fibroblast assembloid models for interrogation of the tumor microenvironment in esophageal adenocarcinoma. Cell Rep. Methods 4, 100909. 10.1016/j.crmeth.2024.100909.

21. Farin, H.F., Mosa, M.H., Ndreshkjana, B., Grebbin, B.M., Ritter, B., Menche, C., Kennel, K.B., Ziegler, P.K., Szabo, L., Bollrath, J., et al. (2023). Colorectal cancer organoid-stroma biobank allows subtype-specific assessment of individualized therapy responses. Cancer Discov. 13, 2192–2211. 10.1158/2159-8290.cd-23-0050.

22. Kabiljo, J., Theophil, A., Homola, J., Renner, A.F., Stürzenbecher, N., Ammon, D., Zirnbauer, R., Stang, S., Tran, L., Laengle, J., et al. (2024). Cancer-associated fibroblasts shape early myeloid cell response to chemotherapy-induced immunogenic signals in next generation tumor organoid cultures. J. Immunother. Cancer 12, e009494. 10.1136/jitc-2024-009494.

23. Dutta, D., Lorenzo-Martín, L.F., Rivest, F., Broguiere, N., Tillard, L., Ragusa, S., Brandenberg, N., Höhnel, S., Saugy, D., Rusakiewicz, S., et al. (2024). Probing the killing potency of tumor-infiltrating lymphocytes on microarrayed colorectal cancer tumoroids. npj Precis. Oncol. 8, 179. 10.1038/s41698-024-00661-3.

24. Dijkstra, K.K., Cattaneo, C.M., Weeber, F., Chalabi, M., van de Haar, J., Fanchi, L.F., Slagter, M., van der Velden, D.L., Kaing, S., Kelderman, S., et al. (2018). Generation of Tumor-Reactive T Cells by Co-culture of Peripheral Blood Lymphocytes and Tumor Organoids. Cell 174, 1586–1598.e12. 10.1016/j.cell.2018.07.009.

25. Recaldin, T., Steinacher, L., Gjeta, B., Harter, M.F., Adam, L., Kromer, K., Mendes, M.P., Bellavista, M., Nikolaev, M., Lazzaroni, G., et al. (2024). Human organoids with an autologous tissue-resident immune compartment. Nature 633, 165–173. 10.1038/s41586-024-07791-5.

26. Teijeira, A., Migueliz, I., Garasa, S., Karanikas, V., Luri, C., Cirella, A., Olivera, I., Cañamero, M., Alvarez, M., Ochoa, M.C., et al. (2022). Three-dimensional colon cancer organoids model the response to CEA-CD3 T-cell engagers. Theranostics 12, 1373–1387. 10.7150/thno.63359.

27. Harter, M.F., Recaldin, T., Gerard, R., Avignon, B., Bollen, Y., Esposito, C., Guja-Jarosz, K., Kromer, K., Filip, A., Aubert, J., et al. (2024). Analysis of off-tumour toxicities of T-cell-engaging bispecific antibodies via donor-matched intestinal organoids and tumouroids. Nat. Biomed. Eng. 8, 345–360. 10.1038/s41551-023-01156-5.

28. Zhang, Y., Hu, Q., Pei, Y., Luo, H., Wang, Z., Xu, X., Zhang, Q., Dai, J., Wang, Q., Fan, Z., et al. (2024). A patient-specific lung cancer assembloid model with heterogeneous tumor microenvironments. Nature Communications 15. 10.1038/s41467-024-47737-z.

29. Jia, Q., Wang, A., Yuan, Y., Zhu, B., and Long, H. (2022). Heterogeneity of the tumor immune microenvironment and its clinical relevance. Experimental Hematology & Oncology 11. 10.1186/s40164-022-00277-y.

30. Polak, R., Zhang, E.T., and Kuo, C.J. (2024). Cancer organoids 2.0: modelling the complexity of the tumour immune microenvironment. Nat. Rev. Cancer 24, 523–539. 10.1038/s41568-024-00706-6.

31. Pelka, K., Hofree, M., Chen, J.H., Sarkizova, S., Pirl, J.D., Jorgji, V., Bejnood, A., Dionne, D., Ge, W.H., Xu, K.H., et al. (2021). Spatially organized multicellular immune hubs in human colorectal cancer. Cell 184, 4734–4752.e20. 10.1016/j.cell.2021.08.003.

32. Rauner, G., Gupta, P.B., and Kuperwasser, C. (2025). From 2D to 3D and beyond: the evolution and impact of in vitro tumor models in cancer research. Nat. Methods 22, 1776–1787. 10.1038/s41592-025-02769-1.

33. Winkler, J., Abisoye-Ogunniyan, A., Metcalf, K.J., and Werb, Z. (2020). Concepts of extracellular matrix remodelling in tumour progression and metastasis. Nature Communications 11. 10.1038/s41467-020-18794-x.

34. Huang, J., Zhang, L., Wan, D., Zhou, L., Zheng, S., Lin, S., and Qiao, Y. (2021). Extracellular matrix and its therapeutic potential for cancer treatment. Signal Transduction and Targeted Therapy 6. 10.1038/s41392-021-00544-0.

35. Asaga, H., Kikuchi, S., and Yoshizato, K. (1991). Collagen gel contraction by fibroblasts requires cellular fibronectin but not plasma fibronectin. Exp. Cell Res. 193, 167–174. 10.1016/0014-4827(91)90552-6.

36. Legant, W.R., Chen, C.S., and Vogel, V. (2012). Force-induced fibronectin assembly and matrix remodeling in a 3D microtissue model of tissue morphogenesis. Integr. Biol. 4, 1164–1174. 10.1039/c2ib20059g.

37 . Fleming, M., Ravula, S., Tatishchev, S.F., and Wang, H.L. (2012). Colorectal carcinoma: Pathologic aspects. J Gastrointest Oncol 3, 153–173. 10.3978/j.issn.2078-6891.2012.030.

38. Blumenthal, R.D., Leon, E., Hansen, H.J., and Goldenberg, D.M. (2007). Expression patterns of CEACAM5 and CEACAM6 in primary and metastatic cancers. BMC Cancer 7. 10.1186/1471-2407-7-2.

39. Eriksen, A.C., Andersen, J.B., Lindebjerg, J., Christensen, R. dePont, Hansen, T.F., Kjær-Frifeldt, S., and Sørensen, F.B. (2018). Does heterogeneity matter in the estimation of tumour budding and tumour stroma ratio in colon cancer? Diagn. Pathol. 13, 20. 10.1186/s13000-018-0697-9.

40. Inoue, H., Kudou, M., Shiozaki, A., Kosuga, T., Shimizu, H., Kiuchi, J., Arita, T., Konishi, H., Komatsu, S., Kuriu, Y., et al. (2023). Value of the Tumor-Stroma Ratio and Structural Heterogeneity Measured by a Novel Semiautomatic Image Analysis Technique for Predicting Survival in Patients With Colon Cancer. Dis. Colon Rectum 66, 1449–1461. 10.1097/dcr.0000000000002570.

41. Fan, P., Zhang, N., Candi, E., Agostini, M., Piacentini, M., Francesca, B., Pierluigi, B., Alessandro, M., Giuseppe, N., Valentina, R., et al. (2023). Alleviating hypoxia to improve cancer immunotherapy. Oncogene 42, 3591–3604. 10.1038/s41388-023-02869-2.

42. Chen, Z., Han, F., Du, Y., Shi, H., and Zhou, W. (2023). Hypoxic microenvironment in cancer: molecular mechanisms and therapeutic interventions. Signal Transduction and Targeted Therapy 8. 10.1038/s41392-023-01332-8.

43. Singleton, D.C., Macann, A., and Wilson, W.R. (2021). Therapeutic targeting of the hypoxic tumour microenvironment. Nature Reviews Clinical Oncology 18, 751–772. 10.1038/s41571-021-00539-4.

44. Lotfi, R., Kaltenmeier, C., Lotze, M.T., and Bergmann, C. (2016). Until Death Do Us Part: Necrosis and Oxidation Promote the Tumor Microenvironment. Transfusion Medicine and Hemotherapy 43, 120–132. 10.1159/000444941.

45. Airley, R., Evans, A., Mobasheri, A., and Hewitt, S.M. (2010). Glucose transporter Glut-1 is detectable in peri-necrotic regions in many human tumor types but not normal tissues: Study using tissue microarrays. Ann. Anat. - Anat. Anz. 192, 133–138. 10.1016/j.aanat.2010.03.001.

46 . Jenkins, W.T., Evans, S.M., and Koch, C.J. (2000). Hypoxia and necrosis in rat 9L glioma and Morris 7777 hepatoma tumors: comparative measurements using EF5 binding and the Eppendorf needle electrode. Int. J. Radiat. Oncol.Biol.Phys. 46, 1005–1017. 10.1016/s0360-3016(99)00342-9.

47. Demmler, R., Anchang, C.G., Yong, Y., Ramming, A., Rauber, S., Schellerer, V.S., Schmid, B., Hartmann, A., Merkel, S., Imkeller, K., et al. (2026). Fibroblast dynamics in colorectal cancer: stability, plasticity, and novel markers. Oncogene 45, 2299–2312. 10.1038/s41388-026-03809-6.

48. Yang, J.-P., Kulkarni, N.N., Yamaji, M., Shiraishi, T., Pham, T., Do, H., Aiello, N., Shaw, M., Nakamura, T., Abiru, A., et al. (2024). Unveiling immune cell response disparities in human primary cancer-associated fibroblasts between two- and three-dimensional cultures. PLOS ONE 19, e0314227. 10.1371/journal.pone.0314227.

49. Strating, E., Verhagen, M.P., Wensink, E., Dünnebach, E., Wijler, L., Aranguren, I., Cruz, A.S.D. la, Peters, N.A., Hageman, J.H., van der Net, M.M.C., et al. (2023). Co-cultures of colon cancer cells and cancer-associated fibroblasts recapitulate the aggressive features of mesenchymal-like colon cancer. Front. Immunol. 14, 1053920. 10.3389/fimmu.2023.1053920.

50. Makutani, Y., Kawakami, H., Tsujikawa, T., Yoshimura, K., Chiba, Y., Ito, A., Kawamura, J., Haratani, K., and Nakagawa, K. (2022). Contribution of MMP14-expressing cancer-associated fibroblasts in the tumor immune microenvironment to progression of colorectal cancer. Front. Oncol. 12, 956270. 10.3389/fonc.2022.956270.

51. Boeck, A.D., Pauwels, P., Hensen, K., Rummens, J.-L., Westbroek, W., Hendrix, A., Maynard, D., Denys, H., Lambein, K., Braems, G., et al. (2013). Bone marrow-derived mesenchymal stem cells promote colorectal cancer progression through paracrine neuregulin 1/HER3 signalling. Gut 62, 550. 10.1136/gutjnl-2011-301393.

52. Gong, Y., Scott, E., Lu, R., Xu, Y., Oh, W.K., and Yu, Q. (2013). TIMP-1 Promotes Accumulation of Cancer Associated Fibroblasts and Cancer Progression. PLoS ONE 8, e77366. 10.1371/journal.pone.0077366.

53. Lu, X., Gou, Z., Chen, H., Li, L., Chen, F., Bao, C., Bu, H., and Zhang, Z. (2025). Extracellular matrix cancer-associated fibroblasts promote stromal fibrosis and immune exclusion in triple-negative breast cancer. J. Pathol. 265, 385–399. 10.1002/path.6395.

54. Lu, G., Hickey, J.W., Haist, M., Qin, X., Zhao, E., Naveed, A., Forgo, E., Baertsch, M.-A., Mani, L., Rovira-Clavé, X., et al. (2026). Single-cell spatial pharmacobiology identifies conserved stromal barriers to therapeutic antibody delivery in human solid tumors. Nat. Biotechnol., 1–15. 10.1038/s41587-026-03152-x.

55. Radtke, F., Clevers, H., and Riccio, O. (2006). From Gut Homeostasis to Cancer. Curr. Mol. Med. 6, 275–289. 10.2174/156652406776894527.

56. Kotliar, D., Veres, A., Nagy, M.A., Tabrizi, S., Hodis, E., Melton, D.A., and Sabeti, P.C. (2019). Identifying gene expression programs of cell-type identity and cellular activity with single-cell RNA-Seq. eLife 8, e43803. 10.7554/elife.43803.

57. Browaeys, R., Saelens, W., and Saeys, Y. (2020). NicheNet: modeling intercellular communication by linking ligands to target genes. Nat. Methods 17, 159–162. 10.1038/s41592-019-0667-5.

58. Jackson, H.W., Fischer, J.R., Zanotelli, V.R.T., Ali, H.R., Mechera, R., Soysal, S.D., Moch, H., Muenst, S., Varga, Z., Weber, W.P., et al. (2020). The single-cell pathology landscape of breast cancer. Nature 578, 615–620. 10.1038/s41586-019-1876-x.

59. Hoch, T., Schulz, D., Eling, N., Gomez, J.M., Levesque, M.P., and Bodenmiller, B. (2022). Multiplexed imaging mass cytometry of the chemokine milieus in melanoma characterizes features of the response to immunotherapy. Sci Immunol 7, eabk1692. 10.1126/sciimmunol.abk1692.

60. Wu, Z.-X., Da, T.-T., Huang, C., Wang, X.-Q., Li, L., Zhao, Z.-B., Yin, T.-T., Ma, H.-Q., Lian, Z.-X., Long, J., et al. (2024). CD69+CD103+CD8+ tissue-resident memory T cells possess stronger anti-tumor activity and predict better prognosis in colorectal cancer. Cell Commun. Signal. 22, 608. 10.1186/s12964-024-01990-3.

61. Boutet, M., Gauthier, L., Leclerc, M., Gros, G., de Montpreville, V., Théret, N., Donnadieu, E., and Mami-Chouaib, F. (2016). TGFβ Signaling Intersects with CD103 Integrin Signaling to Promote T-Lymphocyte Accumulation and Antitumor Activity in the Lung Tumor Microenvironment. Cancer Res. 76, 1757–1769. 10.1158/0008-5472.can-15-1545.

62. Bied, M., Ho, W.W., Ginhoux, F., and Blériot, C. (2023). Roles of macrophages in tumor development: a spatiotemporal perspective. Cellular & Molecular Immunology 20, 983–992. 10.1038/s41423-023-01061-6.

63. Li, N., Zhu, Q., Tian, Y., Ahn, K.J., Wang, X., Cramer, Z., Jou, J., Folkert, I.W., Yu, P., Adams-Tzivelekidis, S., et al. (2023). Mapping and modeling human colorectal carcinoma interactions with the tumor microenvironment. Nat. Commun. 14, 7915. 10.1038/s41467-023-43746-6.

64. Maisel, B.A., Yi, M., Peck, A.R., Sun, Y., Hooke, J.A., Kovatich, A.J., Shriver, C.D., Hu, H., Nevalainen, M.T., Tanaka, T., et al. (2022). Spatial Metrics of Interaction between CD163-Positive Macrophages and Cancer Cells and Progression-Free Survival in Chemo-Treated Breast Cancer. Cancers 14, 308. 10.3390/cancers14020308.

65. Skytthe, M.K., Graversen, J.H., and Moestrup, S.K. (2020). Targeting of CD163+ Macrophages in Inflammatory and Malignant Diseases. International Journal of Molecular Sciences 21, 5497. 10.3390/ijms21155497.

66. Wang, B., Li, Q., Qin, L., Zhao, S., Wang, J., and Chen, X. (2011). Transition of tumor-associated macrophages from MHC class IIhi to MHC class IIlow mediates tumor progression in mice. BMC Immunology 12, 43. 10.1186/1471-2172-12-43.

67 . Buechler, C., Ritter, M., Orsó, E., Langmann, T., Klucken, J., and Schmitz, G. (2000). Regulation of scavenger receptor CD163 expression in human monocytes and macrophages by pro- and antiinflammatory stimuli. J. Leukoc. Biol. 67, 97–103. 10.1002/jlb.67.1.97.

68. Labrijn, A.F., Janmaat, M.L., Reichert, J.M., and Parren, P.W.H.I. (2019). Bispecific antibodies: a mechanistic review of the pipeline. Nature Reviews Drug Discovery 18, 585–608. 10.1038/s41573-019-0028-1.

69. Cioffi, M., Dorado, J., Baeuerle, P.A., and Heeschen, C. (2012). EpCAM/CD3-Bispecific T-cell Engaging Antibody MT110 Eliminates Primary Human Pancreatic Cancer Stem Cells. Clinical Cancer Research 18, 465–474. 10.1158/1078-0432.ccr-11-1270.

70. Bacac, M., Fauti, T., Sam, J., Colombetti, S., Weinzierl, T., Ouaret, D., Bodmer, W., Lehmann, S., Hofer, T., Hosse, R.J., et al. (2016). A Novel Carcinoembryonic Antigen T-Cell Bispecific Antibody (CEA TCB) for the Treatment of Solid Tumors. Clin. Cancer Res. 22, 3286–3297. 10.1158/1078-0432.ccr-15-1696.

71. Friedl, P., and Alexander, S. (2011). Cancer Invasion and the Microenvironment: Plasticity and Reciprocity. Cell 147, 992–1009. 10.1016/j.cell.2011.11.016.

72. García-Rodríguez, J.L., Korsgaard, U., Vissing, S.M., Paasch, T.P., Semenova, M., Vendelbo, S.L., Jensby, E.F., Williams, H.L., Salachan, P.V., Brandt, C.B., et al. (2025). Cancer-associated fibroblasts shape the formation of budding cancer cells at the invasive front of human colorectal cancer. Commun. Biol. 8, 1345. 10.1038/s42003-025-08799-x.

73. Ströhlein, M.A., Siegel, R., Jäger, M., Lindhofer, H., Jauch, K.-W., and Heiss, M.M. (2009). Induction of anti-tumor immunity by trifunctional antibodies in patients with peritoneal carcinomatosis. J. Exp. Clin. Cancer Res. 28, 18. 10.1186/1756-9966-28-18.

74. Segal, N.H., Melero, I., Moreno, V., Steeghs, N., Marabelle, A., Rohrberg, K., Rodriguez-Ruiz, M.E., Eder, J.P., Eng, C., Manji, G.A., et al. (2024). CEA-CD3 bispecific antibody cibisatamab with or without atezolizumab in patients with CEA-positive solid tumours: results of two multi-institutional Phase 1 trials. Nat. Commun. 15, 4091. 10.1038/s41467-024-48479-8.

75. Huynh, P.T., Beswick, E.J., Coronado, Y.A., Johnson, P., O’Connell, M.R., Watts, T., Singh, P., Qiu, S., Morris, K., Powell, D.W., et al. (2016). CD90+ stromal cells are the major source of IL-6, which supports cancer stem-like cells and inflammation in colorectal cancer. Int. J. Cancer 138, 1971–1981. 10.1002/ijc.29939.

76. Leclercq-Cohen, G., Steinhoff, N., Albertí-Servera, L., Nassiri, S., Danilin, S., Piccione, E., Yángüez, E., Hüsser, T., Herter, S., Schmeing, S., et al. (2023). Dissecting the mechanisms underlying the Cytokine Release Syndrome (CRS) mediated by T Cell Bispecific Antibodies. Clin. Cancer Res. 29, 4449–4463. 10.1158/1078-0432.ccr-22-3667.

77. Bengsch, B., Ohtani, T., Herati, R.S., Bovenschen, N., Chang, K.-M., and Wherry, E.J. (2018). Deep immune profiling by mass cytometry links human T and NK cell differentiation and cytotoxic molecule expression patterns. J. Immunol. Methods 453, 3–10. 10.1016/j.jim.2017.03.009.

78. Takata, H., and Takiguchi, M. (2006). Three Memory Subsets of Human CD8+ T Cells Differently Expressing Three Cytolytic Effector Molecules. J. Immunol. 177, 4330–4340. 10.4049/jimmunol.177.7.4330.

79. Forsthuber, A., Aschenbrenner, B., Korosec, A., Jacob, T., Annusver, K., Krajic, N., Kholodniuk, D., Frech, S., Zhu, S., Purkhauser, K., et al. (2024). Cancer-associated fibroblast subtypes modulate the tumor-immune microenvironment and are associated with skin cancer malignancy. Nature Communications 15. 10.1038/s41467-024-53908-9.

80. Mao, X., Xu, J., Wang, W., Liang, C., Hua, J., Liu, J., Zhang, B., Meng, Q., Yu, X., and Shi, S. (2021). Crosstalk between cancer-associated fibroblasts and immune cells in the tumor microenvironment: new findings and future perspectives. Molecular Cancer 20. 10.1186/s12943-021-01428-1.

81. Blair, A.B., Kim, V.M., Muth, S.T., Saung, M.T., Lokker, N., Blouw, B., Armstrong, T.D., Jaffee, E.M., Tsujikawa, T., Coussens, L.M., et al. (2019). Dissecting the Stromal Signaling and Regulation of Myeloid Cells and Memory Effector T Cells in Pancreatic Cancer. Clinical Cancer Research 25, 5351–5363. 10.1158/1078-0432.ccr-18-4192.

82. Knops, A.M., South, A., Rodeck, U., Martinez-Outschoorn, U., Harshyne, L.A., Johnson, J., Luginbuhl, A.J., and Curry, J.M. (2020). Cancer-Associated Fibroblast Density, Prognostic Characteristics, and Recurrence in Head and Neck Squamous Cell Carcinoma: A Meta-Analysis. Front. Oncol. 10, 565306. 10.3389/fonc.2020.565306.

83. Salmon, H., Franciszkiewicz, K., Damotte, D., Dieu-Nosjean, M.-C., Validire, P., Trautmann, A., Mami- Chouaib, F., and Donnadieu, E. (2012). Matrix architecture defines the preferential localization and migration of T cells into the stroma of human lung tumors. J. Clin. Investig. 122, 899–910. 10.1172/jci45817.

84. Tauriello, D.V.F., Palomo-Ponce, S., Stork, D., Berenguer-Llergo, A., Badia-Ramentol, J., Iglesias, M., Sevillano, M., Ibiza, S., Cañellas, A., Hernando-Momblona, X., et al. (2018). TGFβ drives immune evasion in genetically reconstituted colon cancer metastasis. Nature 554, 538–543. 10.1038/nature25492.

85. Chen, Y., McAndrews, K.M., and Kalluri, R. (2021). Clinical and therapeutic relevance of cancer-associated fibroblasts. Nat. Rev. Clin. Oncol. 18, 792–804. 10.1038/s41571-021-00546-5.

86. Seckinger, A., Majocchi, S., Moine, V., Nouveau, L., Ngoc, H., Daubeuf, B., Ravn, U., Pleche, N., Calloud, S., Broyer, L., et al. (2023). Development and characterization of NILK-2301, a novel CEACAM5xCD3 κλ bispecific antibody for immunotherapy of CEACAM5-expressing cancers. J. Hematol. Oncol. 16, 117. 10.1186/s13045-023-01516-3.

87. Sahai, E., Astsaturov, I., Cukierman, E., DeNardo, D.G., Egeblad, M., Evans, R.M., Fearon, D., Greten, F.R., Hingorani, S.R., Hunter, T., et al. (2020). A framework for advancing our understanding of cancer-associated fibroblasts. Nat. Rev. Cancer 20, 174–186. 10.1038/s41568-019-0238-1.

88. Desbois, M., and Wang, Y. (2021). Cancer-associated fibroblasts: Key players in shaping the tumor immune microenvironment. Immunol. Rev. 302, 241–258. 10.1111/imr.12982.

89 . Trüb, M., Uhlenbrock, F., Claus, C., Herzig, P., Thelen, M., Karanikas, V., Bacac, M., Amann, M., Albrecht, R., Ferrara-Koller, C., et al. (2020). Fibroblast activation protein-targeted-4-1BB ligand agonist amplifies effector functions of intratumoral T cells in human cancer. Journal for ImmunoTherapy of Cancer 8, e000238. 10.1136/jitc-2019-000238.

90. Melero, I., Tanos, T., Bustamante, M., Sanmamed, M.F., Calvo, E., Moreno, I., Moreno, V., Hernandez, T., Garcia, M.M., Rodriguez-Vida, A., et al. (2023). A first-in-human study of the fibroblast activation protein-targeted, 4-1BB agonist RO7122290 in patients with advanced solid tumors. Sci Transl Med 15, eabp9229. 10.1126/scitranslmed.abp9229.

91. Claus, C., Ferrara, C., Xu, W., Sam, J., Lang, S., Uhlenbrock, F., Albrecht, R., Herter, S., Schlenker, R., Hüsser, T., et al. (2019). Tumor-targeted 4-1BB agonists for combination with T cell bispecific antibodies as off-the-shelf therapy. Sci. Transl. Med. 11, eaav5989. 10.1126/scitranslmed.aav5989.

92. Melero, I., Tanos, T., Aller, E.C., Qvortrup, C., van Dongen, M., Baraibar, I., Beom, S.-H., Thistlethwaite, F., Riesco, M. del C., Garcia, M.M., et al. (2026). Cibisatamab and FAP-4-1BBL in microsatellite-stable colorectal cancer: a phase 1b trial. Nat. Med. 32, 2430–2439. 10.1038/s41591-026-04380-z.

93. Junttila, M.R., and Sauvage, F.J. de (2013). Influence of tumour micro-environment heterogeneity on therapeutic response. Nature 501, 346–354. 10.1038/nature12626.

94. Quail, D.F., and Joyce, J.A. (2013). Microenvironmental regulation of tumor progression and metastasis. Nat. Med. 19, 1423–1437. 10.1038/nm.3394.

95. Stadler, M., Pudelko, K., Biermeier, A., Walterskirchen, N., Gaigneaux, A., Weindorfer, C., Harrer, N., Klett, H., Hengstschläger, M., Schüler, J., et al. (2021). Stromal fibroblasts shape the myeloid phenotype in normal colon and colorectal cancer and induce CD163 and CCL2 expression in macrophages. Cancer Lett. 520, 184–200. 10.1016/j.canlet.2021.07.006.

96. Floc’h, A.L., Jalil, A., Vergnon, I., Chansac, B.L.M., Lazar, V., Bismuth, G., Chouaib, S., and Mami-Chouaib, F. (2007). αEβ7 integrin interaction with E-cadherin promotes antitumor CTL activity by triggering lytic granule polarization and exocytosis. J. Exp. Med. 204, 559–570. 10.1084/jem.20061524.

97. Postat, J., Merino, M., Mingarelli, A.R., Cerf, A., Bhagrath, A., Patel, D., Shen, C., Brodbeck, J., Tirgar, P., Rogers, D., et al. (2026). Mechanosensing by T cells promotes a tissue-resident memory transcriptional program. Nat. Immunol. 27, 1708–1721. 10.1038/s41590-026-02581-9.

98. Mackay, L.K., Rahimpour, A., Ma, J.Z., Collins, N., Stock, A.T., Hafon, M.-L., Vega-Ramos, J., Lauzurica, P., Mueller, S.N., Stefanovic, T., et al. (2013). The developmental pathway for CD103+CD8+ tissue-resident memory T cells of skin. Nat. Immunol. 14, 1294–1301. 10.1038/ni.2744.

99. Oyler-Yaniv, A., Oyler-Yaniv, J., Whitlock, B.M., Liu, Z., Germain, R.N., Huse, M., Altan-Bonnet, G., and Krichevsky, O. (2017). A Tunable Diffusion-Consumption Mechanism of Cytokine Propagation Enables Plasticity in Cell-to-Cell Communication in the Immune System. Immunity 46, 609–620. 10.1016/j.immuni.2017.03.011.

100. Centofanti, E., Wang, C., Iyer, S., Krichevsky, O., Oyler-Yaniv, A., and Oyler-Yaniv, J. (2023). The spread of interferon-γ in melanomas is highly spatially confined, driving nongenetic variability in tumor cells. Proc. Natl. Acad. Sci. 120, e2304190120. 10.1073/pnas.2304190120.

101. Biasci, D., Smoragiewicz, M., Connell, C.M., Wang, Z., Gao, Y., Thaventhiran, J.E.D., Basu, B., Magiera, L., Johnson, T.I., Bax, L., et al. (2020). CXCR4 inhibition in human pancreatic and colorectal cancers induces an integrated immune response. Proc. Natl. Acad. Sci. 117, 28960–28970. 10.1073/pnas.2013644117.

102. Lu, L., Gao, Y., Huang, D., Liu, H., Yin, D., Li, M., Zheng, J., Wang, S., Wu, W., Zhao, L., et al. (2023). Targeting integrin α5 in fibroblasts potentiates colorectal cancer response to PD-L1 blockade by affecting extracellular-matrix deposition. J. Immunother. Cancer 11, e007447. 10.1136/jitc-2023-007447.

103. Mariathasan, S., Turley, S.J., Nickles, D., Castiglioni, A., Yuen, K., Wang, Y., III, E.E.K., Koeppen, H., Astarita, J.L., Cubas, R., et al. (2018). TGFβ attenuates tumour response to PD-L1 blockade by contributing to exclusion of T cells. Nature 554, 544–548. 10.1038/nature25501.

104. Koch, C.J. (2002). [1] Measurement of absolute oxygen levels in cells and tissues using oxygen sensors and 2-nitroimidazole EF5. Methods Enzymol. 352, 3–31. 10.1016/s0076-6879(02)52003-6.

105. Harter, M.F., D’Arcangelo, E., Aubert, J., Lavickova, B., Havnar, C., Stoll, B., Cubela, I., Filip, A.M., Gaspa-Toneu, L., Scherer, J., et al. (2026). High-throughput histopathology for complex in vitro models. Cell Rep. Methods, 101419. 10.1016/j.crmeth.2026.101419.

106. Chevrier, S., Crowell, H.L., Zanotelli, V.R.T., Engler, S., Robinson, M.D., and Bodenmiller, B. (2018). Compensation of Signal Spillover in Suspension and Imaging Mass Cytometry. Cell Systems 6, 612–620.e5. 10.1016/j.cels.2018.02.010.

107. Greenwald, N.F., Miller, G., Moen, E., Kong, A., Kagel, A., Dougherty, T., Fullaway, C.C., McIntosh, B.J., Leow, K.X., Schwartz, M.S., et al. (2022). Whole-cell segmentation of tissue images with human-level performance using large-scale data annotation and deep learning. Nature Biotechnology 40, 555–565. 10.1038/s41587-021-01094-0.

108. Amitay, Y., Bussi, Y., Feinstein, B., Bagon, S., Milo, I., and Keren, L. (2023). CellSighter: a neural network to classify cells in highly multiplexed images. Nature Communications 14. 10.1038/s41467-023-40066-7.

109. Korsunsky, I., Millard, N., Fan, J., Slowikowski, K., Zhang, F., Wei, K., Baglaenko, Y., Brenner, M., Loh, P., and Raychaudhuri, S. (2019). Fast, sensitive and accurate integration of single-cell data with Harmony. Nat. Methods 16, 1289–1296. 10.1038/s41592-019-0619-0.

110. Germain, P.-L., Lun, A., Meixide, C.G., Macnair, W., and Robinson, M.D. (2021). Doublet identification in single-cell sequencing data using scDblFinder. F1000Research 10, 979. 10.12688/f1000research.73600.2.

